# SLAMF7 and SLAMF8 receptors shape human plasmacytoid dendritic cell responses to intracellular bacteria

**DOI:** 10.1101/2024.04.25.591064

**Authors:** Joaquín Miguel Pellegrini, Anne Keriel, Laurent Gorvel, Sean Hanniffy, Vilma Arce-Gorvel, Mile Bosilkovski, Javier Solera, Stéphane Méresse, Sylvie Mémet, Jean-Pierre Gorvel

## Abstract

Plasmacytoid dendritic cells (pDC), professional type I interferon (IFN) producing cells, have been implicated in host responses against bacterial infections. However, their role in host defense is debated and the operating molecular mechanisms are unknown. Certain Signaling Lymphocyte Activation Molecule Family (SLAMF) members act as microbial sensors and modulate immune functions in response to infection. Here by analyzing multiple human blood transcriptomic datasets, we report the involvement of SLAMF7 and SLAMF8 in many infectious diseases, with elevated levels associated with type I IFN responses in salmonellosis and brucellosis patients. We further identify SLAMF7 and SLAMF8 as key regulators of human pDC function. Silencing of these receptors hinders pDC maturation and abrogates cytokine production during infection with acute (*Salmonell*a) or chronic (*Brucella*) inflammation-inducing bacteria. Mechanistically, we show that SLAMF7 and SLAMF8 signal through NF-κB, IRF7 and STAT-1, and limit mitochondrial ROS accumulation upon *Salmonella* infection. This SLAMF7/8-dependent control of mitochondrial ROS levels favors bacterial persistence and NF-κB activation. Overall, our results unravel essential shared roles of SLAMF7 and SLAMF8 in finely tuning human pDC responses to intracellular bacterial infections with high diagnosis and therapeutic perspectives.

## Introduction

Intracellular bacteria pose significant challenges to the human immune system. Plasmacytoid dendritic cells (pDC) are immune innate cells renowned for their role in antiviral responses, because of their exceptional capacity to produce huge amounts of type I interferon (IFN) (1, 2). Their contribution to defense against bacteria is controversial. Specific deletion of pDC during *Mycobacterium tuberculosis* infection reduces mycobacterial burden and infection pathogenesis in mice (3), in accordance with a detrimental role of pDC during infection with *Citrobacter rodentium* (4), *Listeria monocytogenes* (5) or *Bordetella pertussis* (6). In contrast, pDC depletion worsens the outcome of *Legionella pneumophila* infection in mice (7) and delays post-acute *Klebsiella* pneumonia recovery (8). Human pDC express receptors that regulate their activation and function (1). Yet, the full spectrum of pDC receptors and mechanisms involved during bacterial infection remain obscure.

The Signaling Lymphocyte Activation Molecule Family (SLAMF) of cell surface glycoproteins comprises nine immunoreceptors widely expressed in myeloid and lymphoid cells (9), which have been implicated in diverse immune processes through homotypic interactions, but SLAMF2 and SLAMF4. Some of these receptors serve as microbial sensors. As such, SLAMF1 controls the function of NK cells (10, 11), macrophages (12, 13), DC (14), neutrophils (15), and HSC (16) in response to infection; SLAMF2 and SLAMF6 also interact with bacterial constituents and regulate downstream inflammation (17–19). While some SLAMF members are expressed in pDC, like SLAMF7 and SLAMF9 (20, 21), their role during infection is still to be unraveled.

Here, we reveal the participation of SLAMF7 (CS1, CRACC, CD319) and SLAMF8 (BLAME, CD353) in a broad spectrum of infectious diseases, showing increased levels in the blood of salmonellosis and brucellosis patients that correlate with a type I IFN response. We identify SLAMF7 and SLAMF8 as receptors that positively regulate human pDC function during infection with intracellular bacteria, driving either an acute inflammatory response or a chronic stage. Altogether, this is the first demonstration of an anti-bacterial response in pDC orchestrated by the SLAMF receptors through the activation of the NF-κB, STAT-1, and IRF7 signaling and the regulation of mitochondrial ROS production.

## Results

### Human blood transcriptomic signatures reveal changes in SLAMF7 and SLAMF8 expression levels during infectious diseases

We first conducted a comprehensive analysis of available data sets of blood transcriptomics from healthy individuals and patients suffering from a wide range of infections (including viral (flu, Covid-19), parasitic (malaria), and bacterial (Lyme disease, tuberculosis, staphylococcal infection, streptococcal pharyngitis, brucellosis, salmonellosis) diseases), and highlighted the putative contribution of each of the nine SLAMF receptor family members, given the known ability of some of them to recognize and respond to pathogens. We identified in these unrelated infections and especially in bacterial diseases a striking difference between healthy donors (HD) and infected individuals in the RNA expression of *SLAMF7* and *SLAMF8* (Figure 1A and Supplemental Figure 1A).

**Figure 1.**
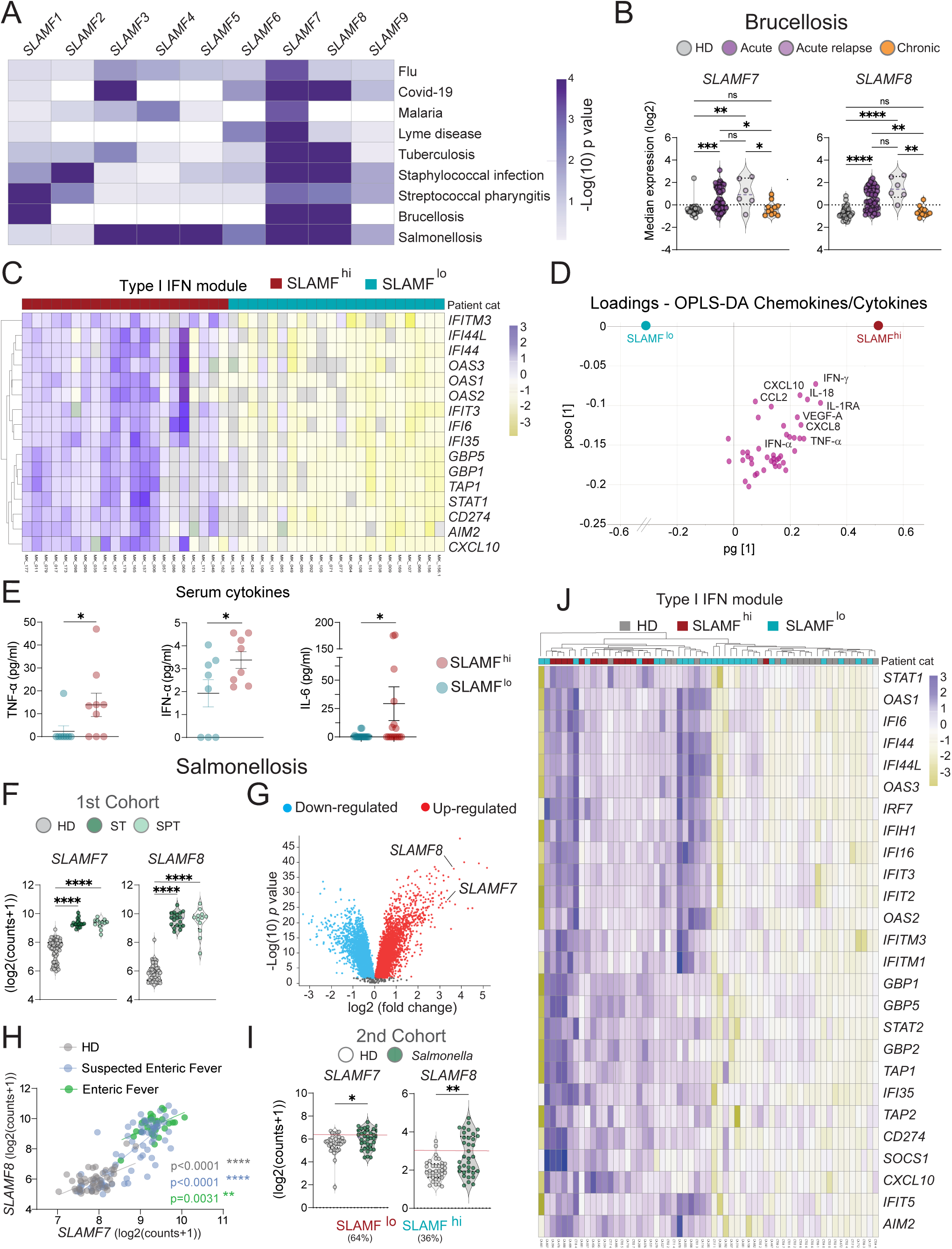
Blood transcriptomic signatures reveal the participation of SLAMF receptors during infection. **A.** Heatmap from blood transcriptomics data showing the differential expression of SLAMF receptors between healthy individuals (HD) and patients suffering from flu (GSE100160), Covid-19 (GSE152075), malaria (GSE116149), Lyme disease (GSE145974), tuberculosis (GSE19491), staphylococcal infection (GSE100165), streptococcal pharyngitis (GSE158163), brucellosis (GSE69597), and salmonellosis (GSE113866). -Log(10) p values are shown with depth of the color representing the degree of variation from HD. **B-C.** RNA seq transcriptomic profiling obtained from whole blood samples from healthy donor (HD) controls or primary brucellosis patients in acute, acute relapse or chronic phase of infection**. B.** The median expression of *SLAMF7* and *SLAMF8* normalized counts is shown. X-axis: HD controls, n=36 (Grey); Brucellosis patients, Acute, n=54 (Purple), Acute with relapse, n=6 (Light Purple), Chronic, n=12 (Orange). Y-axis: log2 residual gene expression counts. Multiple comparison Kruskal-Wallis test, followed by post-hoc Dunn’s test. ns, non-significant. *, p < 0.05; **, p < 0.01; ***, p < 0.001; ****, p < 0.0001. **C.** Heatmap showing type I IFN module gene expression from whole blood RNA-Seq data from SLAMF^hi^ (n=21, Dark Red) and SLAMF^lo^ (n=22, Turquoise) brucellosis patients. **D.** OPLS-DA loading plots of cytokines and chemokines measured in serum, harvested at first visit, from SLAMF^hi^ (n=21, Dark Red) and SLAMF^lo^ (n=22, Turquoise) brucellosis patients. **E.** Univariate analysis of selected cytokine concentration (pg/ml) in serum of SLAMF^hi^ and SLAMF^lo^ brucellosis patients. Significant differences are shown (Mann-Whitney test), *, p < 0.05. **F.** *SLAMF7* and *SLAMF8* normalized counts obtained by transcriptomic profiling analysis from whole blood samples from a first cohort (GSE113866) composed of healthy donor (HD, n=68, Grey) controls, and enteric fever patients with positive cultures for *Salmonella enterica* Typhi (ST, n=19, Dark Green) or *Salmonella enterica* Paratyphi (SPT, n=12, Light Green). Significant differences are shown (Multiple comparison Kruskal-Wallis test, followed by post-hoc Dunn’s test). ****, p < 0.0001, **G.** Volcano plot showing differentially expressed genes between HD and salmonellosis patients from the first cohort (Up-regulated (Red) and Down-regulated (Cyan) in patients). **H.** Correlation between *SLAMF7* and *SLAMF8* RNA counts across all groups of individuals, i. e. HD (Grey), suspected enteric fever (sEF, Pale Blue) and culture-confirmed enteric fever (EF, Pale Green) patients. Non parametric Spearman correlation test. **I.** *SLAMF7* and *SLAMF8* normalized counts obtained by transcriptomic profiling analysis from whole blood samples from a second cohort (GSE69529) composed of HD (n=35, Grey) controls and age-matched patients with *Salmonella* gastroenteritis (n=36, Green). Multiple comparison Kruskal-Wallis test, followed by post-hoc Dunn’s test. *, p < 0.05; **, p < 0.01; ***. **J.** Heatmap showing type I IFN module gene expression from whole blood RNA-Seq data analysis of the second cohort with HD (n=35, Grey) and patients with *Salmonella* gastroenteritis categorized according to their *SLAMF7* and *SLAMF8* expression in SLAMF^hi^ (n=15, Dark Red) and SLAMF^lo^ (n=26, Turquoise) individuals.

### The SLAM-associated signature in brucellosis patients correlates with type I IFN response

We next assessed the participation of SLAMF7 and SLAMF8 in human host defense against *Brucella*, an intracellular bacterium whose peculiarity is to evade the host immune response and establish a chronic infection (22). We analyzed genome-wide transcriptomes that we generated by bulk blood RNA-Seq from HD and brucellosis patients (in acute, acute with relapse or chronic stages at their first visit) (23, 24). Several genes of the SLAM family (*SLAMF1*, *7*, and *8*) were overexpressed in acute but not chronic brucellosis patients, unlike stable *TLR9* gene expression (Figure 1B and Supplemental Figure 1B). Both *SLAMF7* and *SLAMF8* RNA levels were positively correlated in HD and acute brucellosis patients (Supplemental Figure 1C), in spite of the different functions ascribed thus far to SLAMF7 and SLAMF8 (25–30). Discriminative value of SLAMF was investigated by receiver operating characteristic (ROC) curve analysis (Supplemental Figure 1D), giving a ROAUC of 0.7650 (95%CI: 0.6380-0.8919, p=0.0074) for *SLAMF7* and of 0.8736 (95%CI: 0.7764-0.9708, p=0.0002) for *SLAMF8* transcript levels, and confirming the correlation with brucellosis. Categorization of brucellosis patients according to their SLAMF receptor expression indicated a marked difference in the global blood transcriptome of these groups (Supplemental Figure 1E), with *MAPK11* and *VAMP5* as top genes differentially enriched in SLAMF^hi^ patients, and *SESN3* and *MYBPH* in the SLAMF^lo^ group (Supplemental Figure 1F), despite any differences in dominant symptoms and co-morbidities (Supplemental Figure 1G). Importantly, these data unveiled a strong type I IFN signature in SLAMF^hi^ patients as compared with SLAMF^lo^ individuals (Figure 1C). Cytokine and chemokine levels potentially associated with the clinical outcome of *Brucella* infection were measured in the sera of SLAMF^hi^ and SLAMF^lo^ brucellosis patients and analyzed by orthogonal Partial Least Square regression. Cytokine secretion was enriched in sera from SLAMF^hi^ patients, in which top discriminatory analytes included IFN-γ, IL-18, CXCL10, and IL-1RA (Figure 1D). Significantly higher levels of TNF-α, IFN-α, and IL-6 also distinguished SLAMF^hi^ from SLAMF^lo^ brucellosis patients (Figure 1E). Altogether these data identify SLAMF7 and SLAMF8 as hallmarks of acute brucellosis and link their overexpression to type I IFN responses.

### The SLAM-associated signature in salmonellosis patients correlates with type I IFN response

*SLAMF7* and *SLAMF8* genes were differentially expressed in the blood transcriptome of patients suffering from infection with *Salmonella*, a paradigm of acute inflammation-driving bacterium (Figure 1A). We analyzed in depth a first cohort of salmonellosis patients comprising enteric fever patients with positive cultures for both *Salmonella enterica* Typhi and *Salmonella enterica* Paratyphi (31). Significantly higher *SLAMF7* and *SLAMF8* RNA counts were found in whole blood of infected individuals as compared to HD (Figure 1F), which ranked them in the volcano plot among the 50 most differentially expressed genes (Figure 1G). A significant positive correlation between *SLAMF7* and *SLAMF8* RNA counts appeared across all groups of individuals, i. e. HD, culture-confirmed enteric fever patients, and even suspected enteric fever patients (Figure 1H). A second cohort of patients, comprised of children under 10 years of age afflicted with diarrheal diseases (32), confirmed a significant upregulation of *SLAMF7* and *SLAMF8* in those diagnosed with *Salmonella* gastroenteritis (Figure 1I). Applying recursive-partitioning analysis to define the optimal *SLAMF7* and *SLAMF8* expression cutoffs allowed us to categorize these salmonellosis patients into SLAMF^hi^ (n=15, 36%) and SLAMF^lo^ (n=26, 64%) subgroups, with similar demographic and clinical features (Supplemental Figure 1H). Strikingly in both cohorts, *Salmonella*-infected patients showed an elevated type I IFN signature as compared to HD, which was particularly enriched in SLAMF^hi^ patients (Figure 1J and Supplemental Figure 1I-J).

### SLAMF7 and SLAMF8 are overexpressed in human pDC upon TLR7/8 stimulation

Considering that SLAMF receptors are predominantly expressed in innate and adaptative immune cells, we then assessed SLAMF7 and SLAMF8 expression in the discrete immune cell populations of peripheral blood mononuclear cells from HD at steady state and after stimulation with TLR ligands by spectral flow cytometry. pDC were the cell sub-population expressing the highest basal levels of SLAMF7^+^ or SLAMF8^+^ cells (Figure 2A), with an elevated expression close to that of monocytes after stimulation with the ligand of TLR7/8 R848 (Supplemental Figure 2A).

**Figure 2.**
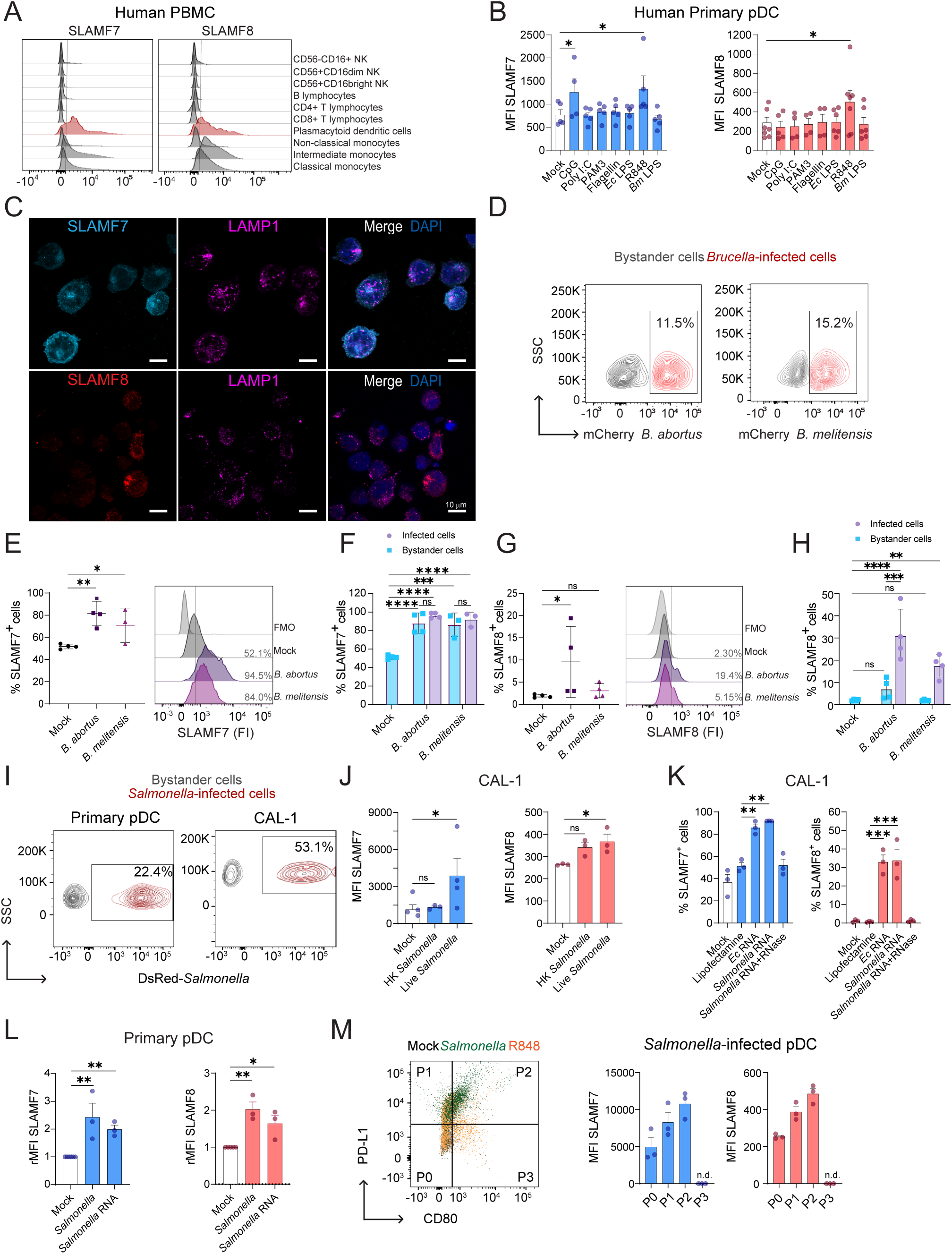
Human pDC infection by *Brucella* or *Salmonella* induces SLAMF7 and SLAMF8 expression. **A.** SLAMF7 and SLAMF8 surface expression on the different cell types of peripheral blood mononuclear cells from healthy donors (n=6) were analyzed by spectral flow cytometry. Representative histograms with positivity indicated by a vertical grey bar are shown. **B.** Purified human primary plasmacytoid dendritic cells (pDC) were stimulated with CpG (TLR9 ligand, 100 ng/mL), Poly I:C (TLR3 ligand, 100 ng/mL), PAM3CSK4 (TLR1/2 ligand, 100 ng/mL), flagellin (TLR5 ligand, 100 ng/mL), *E. coli* LPS (TLR4 ligand, 100 ng/mL), R848 (TLR7/8 ligand, 100 ng/mL), or *B. melitensis* LPS (10 μg/mL) for 24 h. Then, SLAMF7 and SLAMF8 levels of expression were evaluated by flow cytometry. Histograms presenting median fluorescence intensity (MFI) of SLAMF7 and SLAMF8 are shown. Mean ± SD. n=7. Significant differences are indicated. **C.** Representative confocal microscopy images of CAL-1 cells in basal conditions stained for SLAMF7 (Turquoise, top panel), SLAMF8 (Red, bottom panel), LAMP-1 (Magenta) and Nuclei (Blue). Scale bars: 10 μm. n=5. **D.** Representative flow cytometry contour plot graphs showing percentage of CAL-1 cells infected with *Brucella abortus* (left panel) or *Brucella melitensis* (right panel) at 48 h post infection (p.i.) (M.O.I. of 5000) with infected (Red) and bystander (Black) cells indicated. n=4. **E-H.** CAL-1 cells were infected with *B. abortus* (Purple) or *B. melitensis* (Pink purple) (M.O.I. of 5000), and SLAMF7 or SLAMF8 expression was evaluated at 48 h p.i. by flow cytometry. **E.** Left panel, percentage of total SLAMF7^+^ cells is shown with each dot representing an independent experiment. Mean ± SD. n=4. Right panel, representative histograms of flow cytometry experiments showing SLAMF7 expression (fluorescence intensity, FI) with positivity indicated by a vertical grey bar. **F.** Column graphs showing percentages of SLAMF7^+^ infected (Light purple) or bystander (Light blue) cells with each dot representing an independent experiment are shown. Mean ± SD. **G.** Left panel, percentage of total SLAMF8^+^ cells is shown with each dot representing an independent experiment. Mean ± SD. n=4. Right panel, representative histograms of flow cytometry experiments showing SLAMF8 expression (FI) with positivity indicated by a vertical grey bar. **H.** Column graphs showing percentages of SLAMF8^+^ infected (Light purple) or bystander (Light blue) cells are shown. Mean values ± SD. **I.** Representative flow cytometry contour plot graphs showing percentage of primary pDC (left panel) and CAL-1 cells (right panel) infected with DsRed-WT *Salmonella* at 24 h p.i. (M.O.I. of 25) with infected (Red) and bystander (Black) cells indicated. n=12. **J.** Column graphs showing SLAMF7 and SLAMF8 surface expression (MFI) on heat-killed (HK) *Salmonella*-stimulated, live *Salmonella* Typhimurium-infected or mock CAL-1 cells at 24 h p.i., as determined by flow cytometry. Mean ± SD. n=5. **K.** CAL-1 cells were stimulated with highly purified RNA from *Escherichia coli* (Ec RNA), and *Salmonella enterica* Typhimurium (*Salmonella* RNA) in the presence of lipofectamine reagent ± RNase for 24 h, and surface expression of SLAMF7 or SLAMF8 was determined by flow cytometry. Percentage of SLAMF7^+^ (left) and SLAMF8^+^ (right) cells is shown. Mean ± SD. n=3. **L.** Column graphs showing relative SLAMF7 and SLAMF8 surface expression (rMFI) to mock-treated cells put arbitrarily at 1, of human primary pDC stimulated with *Salmonella* RNA or infected with *Salmonella* Typhimurium (M.O.I. of 25) for 24 h. Mean ± SD. n=3. **M.** Left panel, representative flow cytometry dot plot graph showing CD80 and PD-L1 staining in mock-treated (Black), R848-stimulated (Orange) or *Salmonella*-infected primary pDC (Green), and allowing the identification of four subpopulations corresponding to the different quadrants, based on the expression of CD80 and PD-L1. Middle and right panels, column graphs showing SLAMF7 (left) and SLAMF8 (right) surface expression (MFI) in human primary pDC subpopulations after infection with *Salmonella* Typhimurium (M.O.I. of 25) at 24 h p.i.. Mean ± SD. n=3. Statistical differences were all calculated using One-way ANOVA followed by Dunnett’s multiple comparisons test. *, p < 0.05; **, p < 0.01; *** p < 0.001; **** p < 0.0001. ns, non-significant.

The association of a type I IFN signature with SLAMF7 and SLAMF8 upregulation in brucellosis and salmonellosis patients together with their high expression in pDC and the known role of these cells in type I IFN production led us to explore the expression of these two surface receptors in purified human pDC at resting state and after exposure to various stimuli comprising TLR ligands and bacterial components (Figure 2B and Supplemental Figure 2B). 66% and 10% of primary pDC expressed basal levels of SLAMF7 and SLAMF8, respectively (Supplemental Figure 2B). TLR7 stimulation with R848 significantly increased the expression levels of both receptors in human primary pDC, unlike other TLR ligands or bacterial components such as PolyI:C, PAM3, Flagellin, *E. coli* or *Brucella melitensis* LPS (Figure 2B and Supplemental Figure 2B). The TLR9 ligand CpG upregulated the surface expression of SLAMF7 only (Figure 2B and Supplemental Figure 2B). In CAL-1 cells, a human pDC cell line that recapitulates many features of its primary counterparts, SLAMF7 and SLAMF8 were constitutively expressed in 50% and 5% of cells, respectively (Supplemental Figure 2C). Confocal microscopy revealed an intracellular pool of both molecules, located in vesicles negative for the late compartment marker LAMP1 (Figure 2C). As in primary pDC, CpG or R848 triggered a significant rise in the percentage of SLAMF7^+^ CAL-1 cells, while only R848 increased SLAMF8^+^ CAL-1 cell percentage (Supplemental Figure 2C). Neither other TLR ligand nor exposure to outer membrane vesicles (OMV) derived from *B. abortus* WT or lacking the outer membrane protein Omp25 (Supplemental Figure 2C), which we previously demonstrated to interact with SLAMF1 in murine DC (14), modified the proportion of SLAMF7^+^ and SLAMF8^+^ cells in this human pDC cell line. Altogether, we show that the SLAMF7 and SLAMF8 receptors are expressed in human pDC and upregulated by a TLR7/8 agonist.

### Human pDC infection by *Brucella* or *Salmonella* increases SLAMF7 and SLAMF8 expression

To determine if SLAMF7 and SLAMF8 expression may vary in human pDC during brucellosis, we infected CAL-1 cells with *B. abortus* or *B. melitensis*, the two main pathogenic species causing infection in humans. Both *Brucella* species efficiently infected CAL-1 cells (Figure 2D and Supplemental Figure 2D). *B. abortus* or *B. melitensis* infection highly increased SLAMF7 surface expression (Figure 2E) at 48 h post-infection (p.i.) in infected as well as bystander cells (Figure 2F), suggesting that soluble factor(s) mediate SLAMF7 overexpression in non-infected cells. In contrast, only the percentage of SLAMF8^+^ infected cells (and not bystander cells) was significantly augmented by *B. abortus* or *B. melitensis* infection, hereby explaining the slight rise in the total SLAMF8^+^ CAL1 cell percentage at 48 h p.i. (Figure. 2G-H). Altogether, human pDC upregulate via different mechanisms the levels of SLAMF7 and SLAMF8 upon *Brucella* infection.

We next evaluated SLAMF7 and SLAMF8 expression in pDC upon *Salmonella* Typhimurium infection. By using a DsRed-expressing bacteria, we showed by flow cytometry and confocal microscopy that *Salmonella* infects efficiently human primary pDC as well as CAL-1 cells (Figure 2I and Supplemental Figure 2E). Surface expression of SLAMF7 and SLAMF8, but not SLAMF1, significantly increased in CAL-1 cells infected with live *Salmonella*, as well as the percentage of SLAMF7^+^ and SLAMF8^+^ infected cells (Figure 2J and Supplemental Figure 2F-G). The fact that stimulation with heat-killed *Salmonella* had no effect (Figure 2J), opened up a possible involvement of the ligand of TLR7, the viability-associated pathogen-associated molecular pattern (*vita*-PAMP) microbial RNA (33). Indeed, prokaryotic RNA purified from *E. coli* or *S.* Typhimurium increased the percentage of SLAMF7^+^ and SLAMF8^+^ CAL-1 cells in a RNase-dependent manner (Figure 2K), as well as the surface levels of SLAMF7 and SLAMF8 in primary human pDC like in the case of *Salmonella* infection (Figure 2L). This confirms the role of RNA sensing in SLAMF7 and 8 upregulation in human pDC. Notably, both receptors were enriched in the P1 (PD-L1^+^CD80^-^) and P2 (PD-L1^+^CD80^+^) pDC subsets (Figure 2M), specialized in type I IFN production for the former and displaying both innate and adaptive functions for the latter (34). Altogether, we establish that human pDC upregulate the levels of both SLAMF7 and SLAMF8 upon infection with live *Salmonella*, in a process requiring bacterial viability and prokaryotic RNA recognition.

### SLAMF7 and SLAMF8 are required for full activation of *Brucella*-infected pDC

To explore SLAMF7 and SLAMF8 function in human pDC, we generated specific knock-down (KD) CAL-1 cells by lentiviral transduction of short hairpin RNA (shRNA) targeting these receptors (Supplemental Figure 3A). Stable clones were selected based on GFP expression and specific SLAMF receptor silencing as measured by flow cytometry and RT-qPCR (Supplemental Figure 3B-D). The basal level of SLAMF7 was down-regulated in SLAMF8-KD cells compared to that in non-targeting shRNA control (shCTRL) or CAL-1 cells (Supplemental Figure 3C-D). After R848 stimulation, SLAMF8 expression was also significantly reduced in SLAMF7-KD CAL-1 cells; conversely silencing SLAMF8 decreased SLAMF7 surface expression (Supplemental Figure 3D). These observations were confirmed at the protein and mRNA levels upon infection (Supplemental Figure 3C). Functionally, KD of SLAMF7 or SLAMF8 in CAL-1 cells significantly abrogated the increased expression of maturation markers (HLA-DR, CD80, CD86) and of the co-inhibitory molecule, PD-L1, 24h after treatment with R848 (Supplemental Figure 3E).

Then, we asked about the possible role of the pDC SLAMF7 and SLAMF8 receptors in *Brucella* virulence. Silencing SLAMF7 or SLAMF8 significantly reduced the percentage of *B. abortus* and *B. melitensis*-infected CAL-1 cells 48 h p.i. (Figure 3A). CAL-1 cells after *B. abortus* infection harbored, as expected given *Brucella*’s extended immunosuppressive abilities, a moderate level of pDC activation, with a low increase of maturation marker expression and secretion of TNF-α, the only detectable cytokine by multiplex assay (Figure 3B-C and Supplemental Figure 3F-G). Silencing of SLAMF7 or SLAMF8 significantly diminished the proportion of HLA-DR and CD86-expressing pDC, as well as of cells expressing PD-L1 but only in SLAMF8-silenced pDC for the latter (Figure 3B). In addition, *B. abortus*-elicited TNF-α secretion was completely abolished in the SLAMF7- or SLAMF8-KD cells at 48 h p.i. (Figure 3C). Altogether, these results indicate that SLAMF7 and SLAMF8 are required for full activation of human pDC during *Brucella* infection.

**Figure 3.**
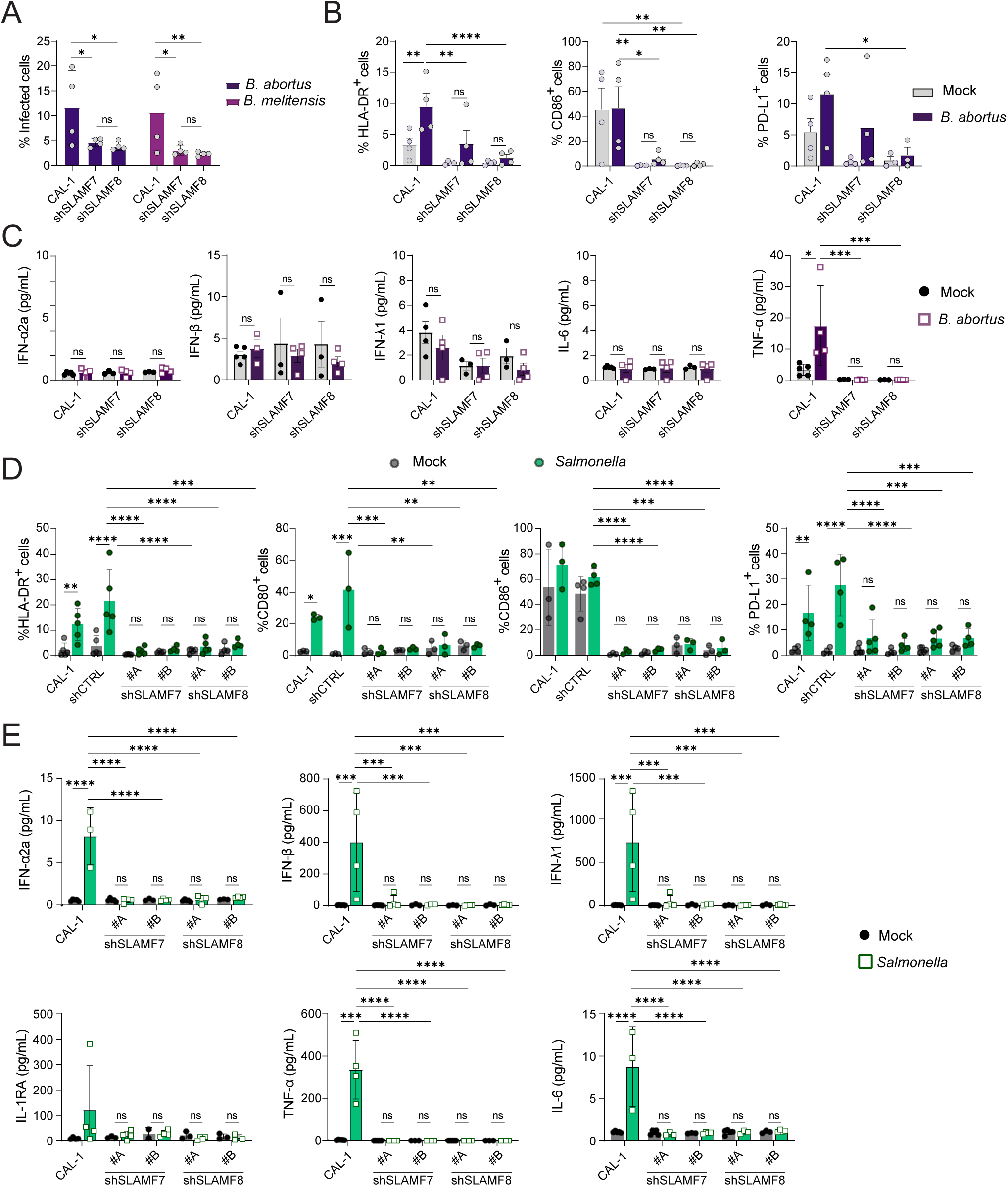
SLAMF7 and SLAMF8 regulate pDC activation and cytokine secretion during intracellular bacterial infection. **A.** CAL-1 and SLAMF-silenced CAL-1 cells were infected with mCherry WT *B. abortus* (Dark purple) or mCherry WT *B. melitensis* (Purple) (M.O.I. of 5000), and cells harboring bacteria were analyzed at 48 h p.i. by flow cytometry. Column graphs showing mCherry *Brucella*^+^ cells. Mean ± SD. n=4. **B-C.** CAL-1 and SLAMF-knocked down (KD) cells were infected with mCherry WT *B. abortus* (M.O.I. of 5000, Dark purple) for 48 h. **B.** Expression of human pDC activation markers was determined by flow cytometry. Column graphs showing percentages of HLA-DR^+^ (left), CD80^+^ (middle) and PD-L1^+^ (right) cells. Mean ± SD. n=4. **C.** Cytokine secretion was determined in culture supernatants using multiplex assay. Column graphs showing cytokine concentration (pg/ml). Each individual point represents one independent experiment. Mean ± SD. n=4. **D-E.** CAL-1, SLAMF-stably transduced CAL-1 cells (2 independent clones, A and B, analyzed per type of SLAM-KD cells, SLAMF7 or SLAMF8, as well as 1 control clone stably transduced with a Shempty vector, shCTRL) were infected with *Salmonella* Typhimurium (M.O.I. of 25, Green) for 24 h. **D.** Expression of activation markers was determined by flow cytometry. Column graphs showing percentages of HLA-DR^+^ (left), CD80^+^ (left center), CD80^+^ (right center) and PD-L1^+^ (right) cells. Each individual point represents one independent experiment. Mean ± SD. n=3 but 5 for HLA-DR. **E.** Cytokine secretion was determined in culture supernatants using multiplex assay. Column graphs showing cytokine concentration (pg/ml). Each individual point represents one independent experiment. Mean ± SD. n=3-4. Statistical differences were all calculated using Two-way ANOVA followed by Sidak’s multiple comparisons test or the nonparametric Mann-Whitney test for unpaired samples. *, p < 0.05; **, p < 0.01; ***, p < 0.001, ****; p < 0.0001. ns. non-significant.

### SLAMF7 and SLAMF8 are essential for activation and cytokine secretion of *Salmonella*- infected human pDC

To determine if these two SLAMF receptors affect human pDC function during *Salmonella* infection, we analyzed our KD and shCTRL clones and CAL-1 cells at 24 h p.i.. Silencing of SLAMF7 and SLAMF8 hindered the maturation of human pDC infected with *Salmonella* when compared to that of shCTRL or CAL-1 cells (Figure 3D), without affecting cell death (Supplemental Figure 3H). Consistently, cytokine secretion by CAL-1 cells, measured by multiplex assay, of type I and III IFN (IFN-α2, IFN-β, IFN-λ1), IL-1RA, TNF-α, and IL-6 was completely abolished in SLAMF7- or SLAMF8-KD cells (Figure 3E). No IFN-γ, IL-1β nor IL-10 production was detected at this time-point whatever the cell type (Supplemental Figure 3I). The impaired activation in SLAMF7- or SLAMF8-KD cells was also evidenced at the RNA level, with significantly reduced expression levels of inflammatory cytokine genes like *IFNB* and *TNFA*, of an IFN-stimulated gene (*ISG15*) and of the gene encoding IκBα, *NFKBIA* (Supplemental Figure 3J). Conversely, the mRNA levels of the anti-inflammatory cytokine IL-10 were greatly elevated (> 500-fold) upon silencing of SLAMF7 or SLAMF8, as compared to CAL-1 or shCTRL cells (Supplemental Figure 3J). Collectively, these findings indicate that SLAMF7 and SLAMF8 play a positive regulatory role in human pDC response to bacterial infection by promoting cell activation and pro-inflammatory cytokine and IFN secretion.

### SLAMF7 and SLAMF8 facilitate *Salmonella* persistence in human pDC by limiting mitochondrial ROS accumulation

Phagocytosis and elimination of hematopoietic tumor cells by macrophages require SLAMF7 independently of SLAM-associated adaptors (35). Hence, we asked if SLAMF7 or SLAMF8 receptors contribute to bacteria phagocytosis by human pDC. Evaluation of bacteria internalization at 2 h p.i, in contrast to later time-points, disclosed no differences in the percentage of DsRed-*Salmonella*^+^ cells between CAL-1 and SLAMF-KD cells (Figure 4A), and in CFU numbers (Figure 4B), consequently excluding a role for SLAMF7 or SLAMF8 in bacteria phagocytosis by pDC. Kinetics experiments showed that intracellular bacterial loads decreased over time in CAL-1 cells, suggesting that pDC actively kill internalized *Salmonella* (Figure 4C). However, SLAMF7- or SLAMF8-KD cells eliminated bacteria more efficiently than CAL-1 and shCTRL cells, as evidenced by a reduced percentage of Ds-Red *Salmonella*^+^ cells, lower CFU and number of bacteria per cell at 24 h p.i. (Figure 4A-B and D). This infers that SLAMF7 and SLAMF8 foster *Salmonella* persistence in pDC.

**Figure 4.**
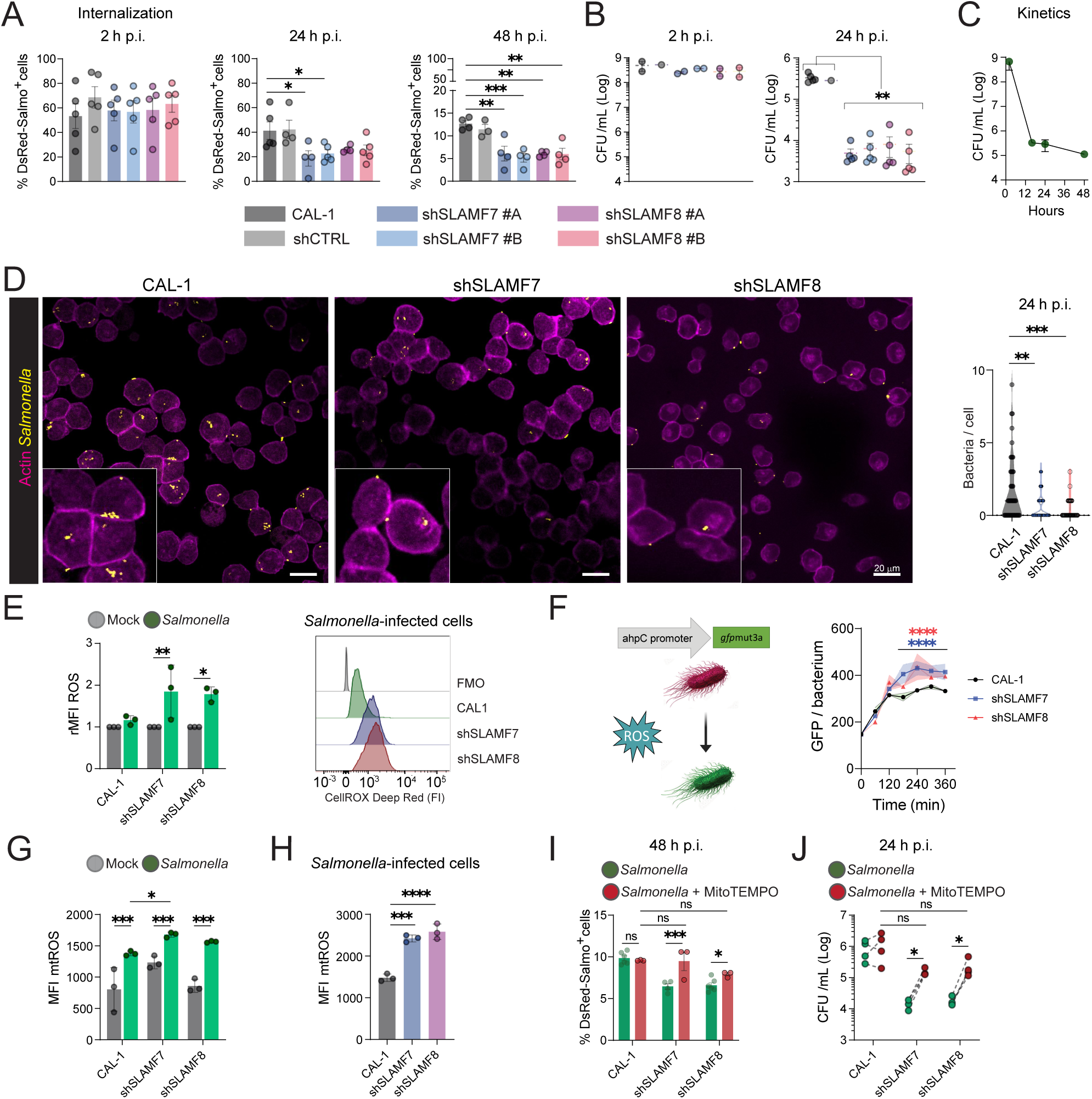
SLAMF7 and SLAMF8 restrict *Salmonella* elimination by pDC by limiting mitochondrial ROS accumulation. **A-C.** CAL-1, SLAMF7- or SLAMF8-silenced (2 clones A and B per SLAMF-KD type) and shCTRL CAL-1 cells were infected with DsRed WT *S.* Typhimurium (M.O.I. of 25) for the indicated times p.i.. One-way ANOVA followed by Dunnett’s multiple comparisons test. **A.** Cells harboring bacteria were analyzed by flow cytometry at 2 h p.i. (Internalization) as well as at 24 h and 48 h p.i.. Column graphs showing percentages of DsRed *Salmonella*^+^ cells. Each individual point represents one independent experiment. Mean ± SD. n=4-5. **B.** Bacterial intracellular burden in infected cells at indicated time-points. Data are represented as CFU/mL (Log), with each individual point representing one independent experiment. Mean ± SD. n=2 (2 h); n=5 (24 h). **C.** Kinetics experiment. *Salmonella* loads (CFU/mL, Log) after infection of CAL-1 cells, and cell incubation until indicated time-points. Each individual point represents one independent experiment. Mean ± SD. n=4. **D.** Left, representative confocal images of CAL-1 and SLAMF-silenced cells infected with DsRed WT *S.* Typhimurium at 24 h p.i.. *Salmonella* (Yellow) and Actin (Purple) are shown. Scale bars, 20 μm. n=3. Right, violin plots depicting the number of bacteria per cell. Pooled data. Multiple comparison Kruskal Wallis test, followed by post-hoc Dunn’s test. **E.** General cellular reactive oxygen species (ROS) in CAL-1 and SLAMF7-or SLAMF8-KD cells loaded with CellROX deep red fluorogenic probe, infected with *S.* Thyphimurium (Green) for 4 h and analyzed by flow cytometry. Left, column graphs of the relative MFI of CellROX deep red (rMFI) to mock-treated CAL-1 (Grey) cells put arbitrarily at 1, with each individual point representing one independent experiment. Mean ± SD. n=3. Right, representative histograms for total ROS content upon *Salmonella* infection are shown (FMO, Fluorescence minus one negative control). **F.** Oxidative stress detection by intracellular *Salmonella*. CAL-1 and SLAMF-silenced cells were infected with the ROS sensing mutant *Salmonella* strain carrying the ahpC-gfp (left scheme). Cells were lysed at different time-points p.i., and GFP fluorescence intensity of intracellular bacteria was determined by flow cytometry. Results presented in the right panel are representative of n=3. **G-H.** Specific mitochondrial superoxide detection in CAL-1 and SLAMF-silenced cells infected with *S.* Thyphimurium for 4 h using MitoSOX and flow cytometry. **G.** Column graphs show MitoSOX MFI in the total population. Each individual point represents one independent experiment. Mean ± SD. n=3. **H.** mtROS levels analyzed in gated *Salmonella*-infected cells from G. **I-J.** After internalization (2 h p.i.), DsRed WT *Salmonella*-infected cells were treated or not with MitoTEMPO (100 µM) and further incubated for the indicated times. **I.** Proportion of DsRed *Salmonella*-infected cells determined at 48 h p.i. by flow cytometry. Each individual point represents one independent experiment. n=4 for CAL-1 and n=3 for SLAMF-KD cells. **J.** Intracellular bacterial burden determined by CFU quantification at 24 h p.i.. n=3. **E-J.** Two-way ANOVA followed by Sidak’s multiple comparisons test. *, p < 0.05; **, p < 0.01; ***, p < 0.001; ****, p < 0.0001.

Given that SLAMF8 inhibits superoxide production by NOX2 complex in murine macrophages (28), and that enhanced activation and oxidative burst reported in SLAMF8-deficient macrophages result in augmented *Salmonella* clearance (26), we analyzed the content of total reactive oxygen species (ROS) in CAL-1 and SLAMF7- or SLAMF8-KD cells first loaded with the CellROX deep red fluorogenic probe and then infected with *Salmonella*. Bacterial infection triggered higher levels of ROS in SLAMF7- or SLAMF8-KD cells as compared to CAL-1 cells (Figure 4E). Then, to monitor the oxidative stress directly sensed by the bacteria, we used a *Salmonella* mutant strain, in which GFP synthesis is upregulated in the presence of exogenous or endogenous H_2_0_2_ (36). As shown in Figure 4F, starting from 180 min p.i. onward, GFP fluorescence was significantly more elevated in bacteria infecting SLAMF7-KD or SLAMF8-KD cells than that observed in CAL-1-infecting bacteria. Silencing SLAMF7 or SLAMF8 in pDC thus augments the oxidative stress sensed by the bacteria.

Since mitochondrial ROS (mtROS) influence type I IFN-producing capacity, activation and antigen cross-presentation effectiveness of human pDC (37, 38), and participate as well in microbial clearance in macrophages (39, 40), we next asked if the enhanced oxidative stress in SLAMF7- or SLAMF8-KD cells involves mtROS. Specific mitochondrial superoxide detection using MitoSOX and flow cytometry revealed that *Salmonella* infection increased mtROS levels in pDC (Figure. 4G), and a significantly higher mtROS accumulation in SLAMF7- or SLAMF8-KD cells as compared to CAL-1 cells (Figure 4H). When inhibiting mtROS in *Salmonella*-infected cells using MitoTEMPO, a mitochondria-targeted antioxidant, percentage of infected CAL-1 cells and intracellular bacterial burden remained stable (Figure 4I), but in SLAMF-KD cells a partial reversion in terms of proportion of DsRed-*Salmonella*^+^ cells and CFU counts occurred (Figure 4J). These results indicate that in human pDC, SLAMF7 and SLAMF8 help *Salmonella* persistence by limiting excessive mtROS accumulation.

### SLAMF7 and SLAMF8 stimulate the NF-κB, IRF7 and STAT-1 pathways in pDC

To decipher the intracellular signaling pathways triggered downstream SLAMF7 and SLAMF8 in human pDC, we performed phospho-flow cytometry on infected cells. Phosphorylation levels of NF-κB p65, IRF7, and STAT1 were significantly higher in *Salmonella* infected CAL-1 or shCTRL cells than those in SLAMF7- or SLAMF8-KD cells (Figure 5A-B). A similar trend was also observed for phosphorylated AKT and p38. Consistently, by confocal microscopy, percentage of phosphorylated NF-κB p65^+^ cells at 3h p.i. was lower when SLAMF7 or SLAMF8 were silenced (Figure 5C). SLAMF receptor functions are controlled by the SLAM-associated protein (SAP) family of adaptors, comprising SAP, EAT-2, and ERT, which either block the recruitment of inhibitory phosphatases (like SHP-1 or SHP-2) or directly activate signaling pathways. These interactions dictate the activation or inhibitory nature of the SLAMF receptors upon engagement (41, 42). Here, we found that EAT-2 was highly expressed in CAL-1 cells, stable upon infection with *Salmonella* and when SLAMF7 or SLAMF8 were down-regulated (Figure 5D). Finally, we examined whether the higher levels of mtROS found in the absence of SLAMF7 or SLAMF8 influence the intracellular signaling initiated in pDC by *Salmonella* infection. Treatment with MitoTEMPO restored the phosphorylation levels of NF-κB p65 in SLAMF7- or SLAMF8-KD cells at 3 h p.i. to those found in CAL-1 or shCTRL cells (Figure 5E). In contrast, mtROS pharmacological inhibition did not affect the phosphorylation of STAT-1 or IRF7 (Figure 5E). These data infer that, in *Salmonella*-infected pDC, moderation of mtROS levels by SLAMF7 or SLAMF8 supports NF-κB activation.

**Figure 5.**
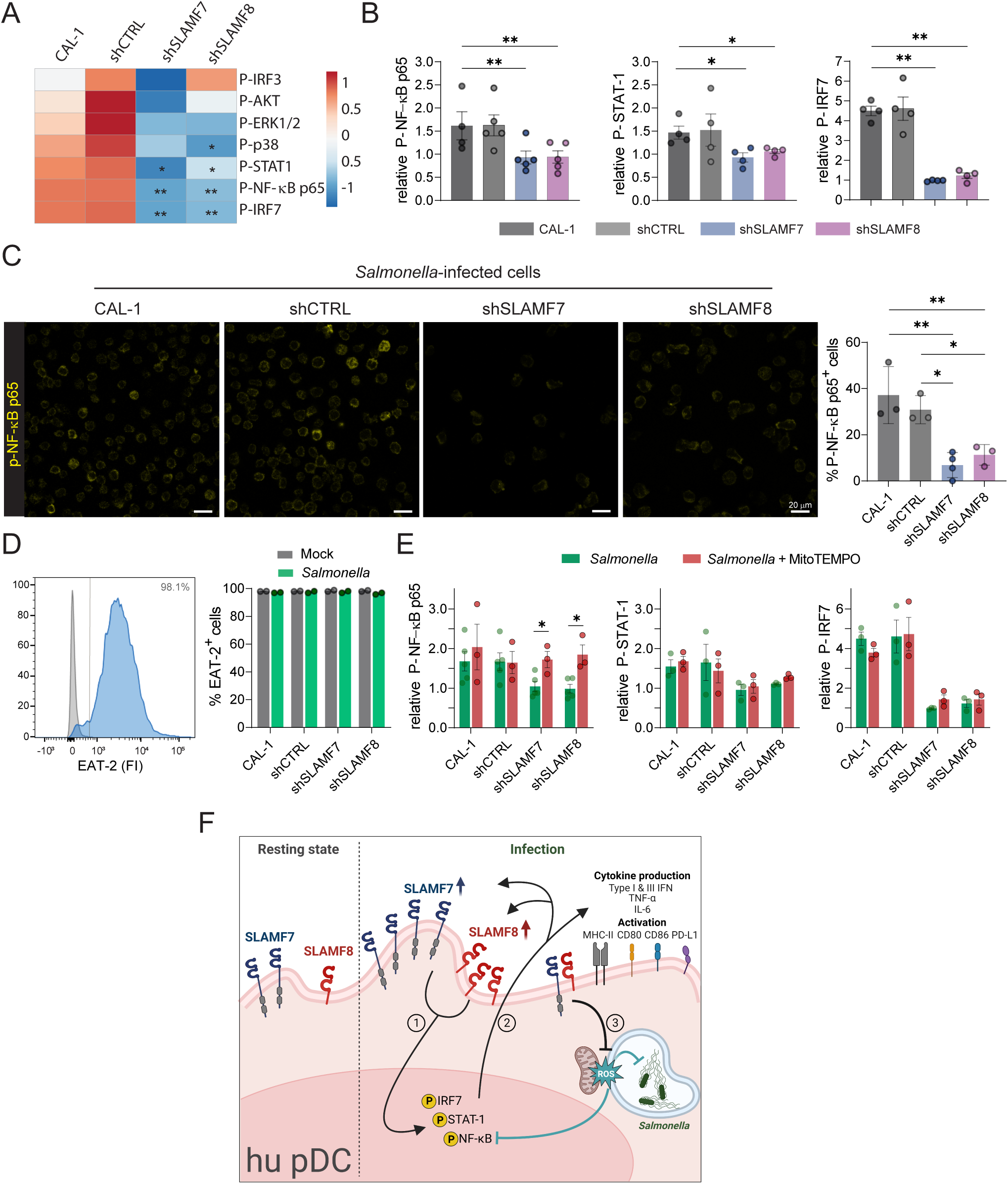
NF-κB, IRF7 and STAT-1 signaling is curtailed in SLAMF7 or SLAMF8 knock-down- pDC. **A-D**. CAL-1 and stably transduced (ShCTRL, shSLAMF7, ShSLAMF8) CAL-1 cells were infected with DsRed WT *S.* Typhimurium (M.O.I. of 25) for 3 h. Then, cells were fixed, permeabilized, and stained for Phospho-flow cytometry or confocal microscopy. **A.** Heatmap showing MFI levels for P-IRF3, P-NF- κB p65, P-IRF7, P-ERK-1/2, P-p38, P-STAT1 and P-AKT of *Salmonella*-infected cells at 3 h p.i. relative to Mock condition. Multiple comparison Kruskal-Wallis test, followed by post-hoc Dunn’s test. n=4. **B.** Column graphs showing relative MFI of phosphorylated proteins (left, P-NF-κB p65; middle, P-STAT-1; right, P-IRF7) in infected cells. Mean ± SD. n=5. **C.** Left, representative confocal images of P-NF-κB p65 in CAL-1 and stably transduced CAL-1 cells infected with *Salmonella*. Scale bars: 20 μm. Right, quantification of P-NF-κB p65^+^ cells. n=3. One-way ANOVA followed by Dunnett’s multiple comparison test. **D.** Left, representative flow cytometry histogram of EAT-2 adaptor expression (fluorescence intensity (FI) in CAL-1 cells with positivity indicated by a vertical grey bar. Right, percentages of EAT- 2^+^ CAL-1 and stably transduced cells infected (Green) or not (Grey) with *S.* Typhimurium. n=3. **E.** CAL- 1 and stably transduced CAL-1 cells were infected with DsRed WT *Salmonella* Typhimurium (M.O.I. of 25) for 1 h. After washes and resuspension in gentamicin-containing medium, cells were treated with MitoTEMPO (100µM) for 2 h. Phospho-flow cytometry was performed to analyze the phosphorylation levels of selected proteins. Column graphs show relative MFI of P-NF-κB p65 (left), P-STAT-1 (middle) and P-IRF7 (right). Mean ± SD. n=4. Two-way ANOVA followed by Sidak’s multiple comparison test. Only statistical differences are shown. *, p < 0.05; **, p < 0.01; *** p < 0.001; **** p < 0.0001. no p value, non-significant. **F.** Proposed model for SLAMF7 and SLAMF8 function in human pDC during intracellular bacterial infection. At the resting state, few SLAMF8 is expressed on the surface of human pDC in contrast to higher SLAMF7 levels. Infection with an intracellular bacterium, like *Salmonella*, engages SLAMF7 and SLAMF8 receptors, and elicits the phosphorylation and activation of IRF7, STAT- 1 and NF-κB in a SLAMF7/8-dependent manner (1). This induces the expression of SLAMF7, SLAMF8, pDC activation markers, as well as cytokine secretion (TNF-α, IL-6 and type I and III IFN) (2). Elevated levels of SLAMF7 or SLAMF8 restrain mitochondrial ROS production, and subsequently favor NF-κB activation as well as *Salmonella* persistence (3). This illustration was created using Biorender.

## Discussion

In this work, we resolve the long-standing debate regarding the direct infectibility and activation of human pDC by bacterial pathogens (43–47). Using two different models of intracellular bacteria leading to acute (*S.* Typhimurium) or chronic (*Brucella* spp.) infection, we demonstrate that these bacteria not only infect pDC but also control pDC function in a SLAMF7 and SLAMF8 co-dependent fashion. We found that in pDC SLAMF7 and SLAMF8 are co-regulated at the protein and mRNA levels in steady state and upon infection or TLR7/8 stimulation. This co-regulation is consistent with the simultaneous variation of *SLAMF7* and *SLAMF8* mRNA expression in our human blood transcriptomic analyses from a wide range of infectious disease patients, but differs from the sole upregulation of SLAMF7, and not SLAMF8, in PBMC and monocytes/macrophages of sepsis patients (48). Using heat-killed *Salmonella* or purified bacterial RNA, we show that SLAMF7 and SLAMF8 upregulation in pDC likely depends on TLR7/8.

Our findings identify human SLAMF7 and SLAMF8 as immunoreceptors that trigger human pDC maturation, activation, pro-inflammatory cytokine and type I IFN secretion, and promote the survival of intracellular bacteria during infection. This activating function of SLAMF7 and 8 during *Brucella* infection in pDC contrasts with the restriction of DC activation driven by the interaction of SLAMF1 with *Brucella* Omp25, which also facilitates bacterium intracellular survival (14) and the negative regulation of pro-inflammatory cytokine production in macrophages by SLAMF7 upon TLR4 stimulation (48). Beside type I and III IFN, we found that *Salmonella*- and *Brucella*-infected pDC secrete TNF-α in a SLAMF7/8-dependent manner, which is coherent with the SLAMF7/8-dependent activation of the NF-κB, STAT-1 and IRF7 pathways that we observed in *Salmonella*-infected cells. These findings concur with the rapid induction of TNF-α upon SLAMF7 engagement on macrophages, amplifying cell activation through an autocrine loop (25), and with the cell-intrinsic TNF signaling that promotes pDC IFN production during MCMV infection *in vivo* (49).

As regards the SAP family of adaptors in human pDC, we found that EAT-2 was stably expressed in *Salmonella*-infected cells in both WT or KD CAL-1 cells. This suggests that in human pDC EAT-2 may interact with SLAMF7 intracellular tyrosine-based switch motifs (ITSM) upon infection, evoking the direct SAP-independent association of SLAMF7 with EAT- 2 that triggers NK cell activation and cytotoxicity (50–52). It remains possible that like for tumor cell elimination by macrophages (35), SLAMF7 downstream signaling during infection occurs without SAP adaptor recruitment. SLAMF8, like SLAMF9, lacks ITSM on its cytoplasmatic tail but can also elicit specific transduction pathways (26, 28, 53) via the interaction with phosphatases, such as SHP-2 (53, 54). Further investigation is needed to unravel pDC SLAMF8 interacting partners, which might also associate with SLAMF7 given their shared behavior in these cells upon infection.

In human pDC, we observed that SLAMF7 or SLAMF8 silencing enhances *Salmonella* clearance, ROS production and the oxidative stress directly sensed by the bacteria, in agreement with prior work on SLAMF8-deficient mouse macrophages (26). We further showed that the higher levels of mtROS seen in SLAMF7- or SLAMF8-KD pDC are responsible for this bacterial clearance. This demonstrates that SLAMF7 and SLAMF8 control mtROS levels to promote bacteria intracellular survival. The tempering of mtROS is most probably achieved upon engagement of SLAMF7 and SLAMF8 through the PI3K pathway since we observed less AKT phosphorylation in SLAMF7/8 KD cells. This in accordance with the reported role of SLAMF8 in macrophages (26). We also found that when SLAMF7 or SLAMF8 is silenced, NF-κB phosphorylation is decreased, but restored to WT levels by the inhibition of mtROS. These findings are consistent with the complex role played by the oxidative stress on NF-κB activation (55) and illustrate the fine tuning of human pDC inflammatory response to infection by SLAMF7 or 8. Overall, while mtROS is known to influence the activation and type I IFN-producing capacity of pDC (37, 38), this work is the first to describe a limitation of mtROS signaling pathways in pDC.

Thus, beside SLAMF9 involved in pDC homeostasis in a full organism (20), SLAMF7 and SLAMF8 emerge as paramount members of the SLAMF receptor family regulating pDC response to intracellular bacteria. The involvement of SLAMF7 and SLAMF8 extends beyond infectious diseases to cancer and autoimmunity (9, 56). Therefore, the concerted role of SLAMF7 and SLAMF8 in orchestrating human pDC responses against bacteria (Figure 5F) suggests that cell-specific blockade of these receptors may constitute a promising therapy to abate pDC activation and hyper-production of type I IFN during infection, as well as in autoimmune disorders and cancer.

## Materials and Methods

### Ethics

The brucellosis study in humans was approved by the Ethics Committee of the Medical Faculty in Skopje, Republic of North Macedonia (authorization 03-7670/2). Laboratory data were obtained during routine diagnostic procedures, and all patients and healthy donors (HD) enrolled gave their written consent.

### Cell culture

Healthy human blood and Buffy coats were obtained by leukapheresis (Etablissement Français du Sang (EFS), Marseille, France). Peripheral blood mononuclear cells (PBMC) were isolated by centrifugation over Ficoll-Hypaque (#17144003 Cytiva Sigma-Aldrich) and cultured (1x10^6^ cells/mL) in flat-bottom 48-well plates with RPMI1640 (#21875034 Gibco Thermofisher) supplemented with L-glutamine (#25030024 Gibco Thermofisher), and 10% fetal bovine serum (FBS) (#P30-3306 PAN BIOTECH Dutscher). Purification of primary pDC was performed using the Plasmacytoid Dendritic Cell Isolation Kit II (#130-097-415 Miltenyi Biotec), reaching >93% of purity. Cells were then infected or stimulated with bacterial components or TLR ligands as specified in the figure legend and analyzed by flow cytometry. RNA stimulation was performed in the presence of lipofectamine to facilitate access to endosomal sensors and protect RNA from degradation.

CAL-1 cells (kind gift from Dr. P. Pierre, CIML, France) were grown in RPMI1640, 10% FBS, 2mM L-glutamine, 1x non-essential amino acids (#11140050 Gibco Thermofisher), 10mM HEPES (#15630080 Gibco Thermofisher), 1 mM sodium pyruvate (#11360039 Gibco Thermofisher). For performing experiments, cells were plated in 24-well plates at 4x10^5^ cells/ml, 1mL/well, in complete medium with 2% FBS. All cell lines were mycoplasma-free as determined with the MycoAlert Mycoplasma Detection Kit (#LT07-318 Lonza). When specified, MitoTEMPO (100 µM, # SML0737 Sigma-Aldrich) was added to cell culture.

### Mutant CAL-1 cells generation

Specific knockdown CAL-1 cells were generated by transduction of lentiviral particles containing short hairpin RNAs (shRNAs) targeting SLAMF7 and SLAMF8. Briefly, specific shRNA-expressing plasmids (pGFP-C-shLentiSLAMF7 and pGFP-C-shLentiSLAMF7, Origene), with pCMV-VSV-G envelope plasmid and pCMV-dR8.91 Gag and Polymerase expression plasmid (kind gifts of M. Negroni, IBMC Strasbourg, France) were transfected into HEK293T cells by calcium phosphate method. 2 days after transfection, the viral supernatant was harvested, filtered through 0.45 µm filter and concentrated on Amicon Ultra-15 centrifugal filters (#C7715 Merck). The concentrated viral particles were added to CAL-1 cells together with lentiBLAST Premium transduction enhancer (1µL, # LBPX500 OZ Biosciences) and centrifuged for 90 min at 800 g. Cells were incubated for 48 h, after which the media was replaced with fresh complete media. Puromycin selection (0.5 µg/mL, #A1113803 Therrmofisher) was then applied to select for cells that had stably integrated the lentiviral construct. Monoclonal cell lines were obtained by limiting dilution. The selected cells were further characterized by flow cytometry and qPCR to confirm knock-down efficiency and GFP expression. Control cells were generated in parallel using lentiviral particles containing non-targeting shRNA. Mutant CAL-1 clones were maintained in complete medium supplemented with puromycin (0.2 µg/mL).

### Bacterial strains and infection

*Salmonella* Typhimurium and isogenic strains used in this study are the following: wild-type (WT) *Salmonella enterica* subsp. *enterica serovar* Typhimurium (*S.* Typhimurium strain 12023, laboratory stock); *S*. Typhimurium DsRed strain (57); *S*. Typhimurium ROS sensing strain, which contains an OxyR-dependent promoter fused to the *gfp*mut3a gene and lacks four H_2_O_2_-degrading enzymes (*katE katG katN ahpCF*) (39). Strains were cultured in Luria Broth medium (Difco) at 37°C and ampicillin (50 μg/ml), kanamycin (50 μg/ml) or chloramphenicol (50 μg/ml) were added when required.

Fluorescent *Brucella* (bIN#1559 for *B. abortus* strain 2308; bIN#1502 for *B. melitensis* strain 16M) were obtained by transformation by a pMR10-based plasmid encoding mCherry (58). They were cultured on Tryptic Soy Agar plates containing kanamycin (25 μg/ml) at 37°C. All *Brucellae* were kept, grown and used under strict biosafety containment conditions all along experiments in the BSL3 facility, of VBIC, Nîmes, France.

*S.* Typhimurium at a multiplicity of infection (M.O.I.) of 25:1 or *Brucella spp.* at a M.O.I. of 5000:1 were used to infect human primary pDC or CAL-1 cells. Bacteria were plated onto cells in 100 µL and left for 1 h at 37 °C with 5% CO_2_. After two washes with medium, cells were incubated for 1 h in medium containing 50 μg/ml gentamicin to kill extracellular bacteria. Thereafter, antibiotics concentration was decreased to 5 μg/ml.

### Flow cytometry

Cells were harvested and stained for 30 min at 4°C with the primary antibody mix. Then, cells were washed in PBS with 2% of FCS and incubated with the secondary antibody mix for 30 min at 4°C. When needed, a third step of antibody incubation was added for staining with coupled antibodies. After washing, cells were stained with LIVE/DEAD Fixable Blue Dead Cell Stain (#L34961 Invitrogen Thermofisher) for 10 min at 20°C. Cells were then fixed for 10 min in 3.2% PFA at 20°C or analyzed immediately. Antibodies used in this study were the following: from Biolegend (SLAMF1 clone A12(7D4) #306302; SLAMF7/CD319/CRACC clone 162.1 #331802; mouse IgG1 clone RMG1-1 #40615; mouse IgG2b κ clone MPC-11 #400342; CD16 clone 3G8, Biolegend, #302026; HLA-DR clone L243 #307640), from Thermofisher: (SLAMF8 clone 250014, #MA523947; PD-L1/CD274 clone MIH1, #46598342; CD274/PD-L1 clone MIH1, # 46-5983-42);; CD303,(clone 201A, eBioscience, #11-9818-42), from BD Biosciences (CD123, clone 9F5, #551065; CD14 clone MDP9 #560180; CD3 clone SK7 #563798, CD56 clone B159, #55551, CD80 clone 307.4 #557227, CD86 clone FUN-1 #563412), from Cytek Biosciences (CD4 clone SK3 #SKU R7-20074; CD8 clone SK1 #SKU R7-20036; CD19 clone SJ25C1 #SKU R7-20274), from Sigma-Aldrich (EAT2/CD81 clone M38, #SAB4700232). Phospho-flow staining was performed by fixing the cells with 3.2% PFA for 10 min at 20°C, followed by permeabilization with cold methanol for 15 min. After washing, intracellular staining with antibodies for phospho-proteins (Phospho-NF-κB p65 (Ser536) clone 93H1, #3033; P-Stat1 (Tyr701) clone D4A7, #7649; P-IRF7 (ser477) clone D7E1W, #42016; Phospho-IRF-3 (Ser396) #29047; P-AKT (thr308) #9275; Phospho-p38 MAPK (Thr180/Tyr182), #9211; Phospho-p44/42 MAPK (Erk1/2) (Thr202/Tyr204) clone D13.14.4E, #4370; all from Cell Signaling Technology) was done in 1xPBS 2%BSA. Flow cytometry was performed using a FACS Aurora (Cytek) for PBMC or a Fortessa (BD Biosciences) for primary and pDC cell lines, and *Salmonella*, and data were analyzed with the Flow-Jo_v9.9.4 software. Surface staining gating relied on the examination of mock cell and isotype control antibody staining.

### Immunofluorescence microscopy

At different time-points of infection, cells were seeded on alcian blue-treated coverslips, fixed with 3.2% PFA, permeabilized with 0.05% saponin, followed by 1 h blocking with 2% BSA in PBS. Primary antibodies (rabbit anti-LAMP1, 1/1000, kind gift of Dr. Minoru Fukuda (La Jolla Cancer Research Foundation, U.S.A.); phalloidin-AF488 for F-actin detection, #A12379 Therrmofisher) were incubated for 1 h in 1xPBS, 0.1% saponin, 0.1% horse serum. After 2 washes in 1xPBS, secondary antibodies were incubated for 45 min, and coverslips were finally mounted in Prolong Gold antifade reagent with DAPI (#P36931 Invitrogen Thermofisher). Images (of 1024x1024 pixels) were acquired on a Leica SP5 laser scanning confocal microscope and assembled using Fiji software V1.53s (ImageJ).

### Bacterial intracellular burden assay

To monitor bacterial intracellular load, infected cells at different time-points were washed 3 times in 1xPBS and lysed with 0.1% Triton X-100 (#X100 Sigma-Aldrich) in H_2_O. Serial dilutions of cell lysates were made and 20 µl aliquots were plated in triplicates onto LB agar. Plates were incubated for 24 h at 37°C, and colonies were counted from dilutions yielding 10 to 100 visible colonies.

### Cytokine measurement

For *Salmonella*- and *Brucella*-infected cells, supernatants were harvested and cytokine measured using Meso Scale Discovery (MSD) U-PLEX Biomarker Group1 (human) Assay (#K15067L-1 Meso Scale Diagnostics) for IFN-α2a, IFN-β, IFN-γ, IL-29/IFN-λ1, TNF-α, IL-10, IL-6, IL-1β, and IL-1RA. MSD assay was performed as per manufacturer’s instructions but with a 16 h incubation of the samples to improve binding to the plate.

For human brucellosis studies, cytokine/chemokine concentrations were determined in serum harvested at initial visit from acute, acute with relapse, chronic infected patient and HD groups by Luminex using the Human Cytokine/Chemokine/Growth Factor 45-Plex ProcartaPlex Panel 1 (#EPX450-12171-901 ThermoFisher) according supplier’s recommendations. The resulting data were normalized and analyzed by OPLS-DA (Orthogonal Partial Least Squares Discriminant Analysis) using the MetaboAnanalyst platform version 5.0.

### ROS detection

To measure ROS, CAL-1 cells were infected with *Salmonella* for 4 h and then incubated for 30 min at 37 °C with CellROX Deep Red Reagent (1 µM, #C10422 Invitrogen Thermofisher) for total (cytoplasmic and nuclear) cellular ROS assessment or with MitoSOX (1,25 µM, #M36009 Invitrogen Thermofisher) in PBS 2%FBS for mtROS detection. Cells were then washed twice with warm PBS, immediately re-suspended in cold PBS containing 1% FBS and subjected to flow cytometry analysis.

To analyze the oxidative stress detected by the bacteria, *Salmonella* WT- and *Salmonella* ROS sensing strain-infected CAL-1 cells were washed twice with PBS, lysed with 0.1% Triton X-100 in PBS and immediately fixed in two volumes of 3% paraformaldehyde for 1 h. Large debris and nuclei were removed by centrifugation for 5 min at 200 g and bacteria were pelleted at 20,000 g for 10 min. Bacteria were resuspended in 40 µL of 10 mM NH_4_Cl in PBS, immunolabelled with a mouse anti-*Salmonella* 1E6 (#MA1-83451 Thermofisher) and a rat anti-mouse IgG1 BV421, both diluted 1:1000 and analyzed by flow cytometry.

### RNA extraction and RT-qPCR

Total RNA was extracted from infected cells using TRIzol (#15596026 Thermofisher) and Direct-zol RNA microprep Kit (#ZR2070T Zymo Research Ozyme) following the manufacturer’s instructions. with an additional on-column DNase RNase-free (#79254 Qiagen) incubation step to get rid of any genomic DNA trace. cDNA were generated with 500 ng of RNA as a template using QuantiTech Reverse Transcription Kit (#205311 Qiagen) according to manufacturer’s recommendations. 2 μl of cDNA corresponding to 6 ng of starting total RNA were used for qPCR with specific primer sets. Amplification reactions were performed in duplicates with SYBR Green mix (#RR420L Takara) in 7500Fast Real-time PCR machine (Applied Biosystems). Gene encoding β-actin, *ACTB*, was selected as the best housekeeping gene in human pDC. Data were normalized with this gene, further analyzed by the ΔΔCt comparative cycle threshold method and presented as relative expression versus Mock-treated cells. Primer sequences were *SLAMF7* forward 5’-TGCCTCACCCTCATCTATATCCT-3’ and reverse 5’- CTTCAGGGGGAAAGTCACGG-3’, *SLAMF8* forward 5’-CTCCGTGTTGATGGTGGACA-3’ and reverse 5’-TGAACACTTGTACCACGGGC-3’, *IFNB* forward 5’-ACCATCTGAAGACAGTCCTGG-3’ and reverse 5’-GTGACTGTACTCCTTGGCCTT-3’, *TNFA*forward 5’-AGTGACAAGCCTGTAGCCCATGTT-3’ and reverse 5’-GTTATCTCTCAGCTCCACGCCATT-3’, *ISG15* forward 5’-GCAGCGAACTCATCTTTGCC-3’ and reverse 5’-TCTTCACCGTCAGGTCCCA-3’, *NFKBIA* forward 5’-CACCGAGGACGGGGACT-3’ and reverse 5’-CACCTGGCGGATCACTTCC-3’, *IL10* forward 5’-CTGGGGGAGAACCTGAAGAC-3’ and reverse 5’-GGCCTTGCTCTTGTTTTCACA-3’, *ACTB* forward 5’-CATGAGAAGTATGACAACAGCC-3’ and reverse 5’-AGTCCTTCCACGATACCAAAGT-3’.

For prokaryotic RNA extraction, 5x10^9^ CFU of *S.* Typhimurium or *E. coli* were resuspended in 1 ml of TRIzol and their RNA was purified using Quick-RNA DirectZol columns ((#ZR2072 Zymo Research Ozyme) according to the manufacturer’s instructions. The quantification and purity of the prokaryotic RNA was determined using a Nanodrop spectrophotometer.

### Transcriptomics analysis

#### Transcriptome dataset selection and population study

We first browsed the Gene Expression Omnibus (GEO) repository using the following keywords and expressions: [(VIRUS) OR (BACTERIA) OR (PARASITE) OR (INFECTION) OR (PARASITE) OR (pathogen)] AND [(transcriptomics) OR (RNA-seq) OR (microarray) OR (expression) OR (transcriptome)] AND [(BLOOD) OR (WHOLE BLOOD)]. All potentially relevant datasets were further evaluated in detail. The eligibility criteria included: (i) publicly available transcriptome data; (ii) detailed sample information; (iii) detailed protocol information; (iv) inclusion of HD controls; (v) same sample source -whole blood-.For salmonellosis detailed human studies, a first cohort of patients was referenced as GSE113866 (31), and a second cohort as GSE69529 (32).

Brucellosis patients were handled at the University Clinic of Infectious Diseases and Febrile Conditions, Skopje, spanning the years 2007 to 2015. The study encompassed a total of 108 patients, categorized as follows: Healthy donors (n=36), Acute treated (n=54), Acute with relapse (n=6), and Chronic (n=12). RNA sequencing was conducted using a protocol established by the Benaroya Research Institute, Seattle, USA. Serum samples from these patients, as well as healthy donor control blood, were stored at -80°C for subsequent cytokine/chemokine analysis. The original RNA-Seq datasets have been deposited in the Gene Expression Omnibus (GEO) repository under accession number GSE69597 (23, 24). The diagnosis of brucellosis was based on clinical signs and symptoms relevant to this disease (fever, sweating, malaise, arthralgia, hepatomegaly, splenomegaly, and focal disease signs), and validated by a qualitative positive Rose Bengal test and a Brucellacapt assay of >1/320.

#### Microarray differential expression analysis

Data were log2-transformed and Force normalization was applied. GEOquery and Limma (Linear Models for Microarray Analysis) were used for differential expression analysis in R. Benjamini & Hochberg false discovery rate method was used for multiple-testing corrections.

#### RNA-Seq differential expression analysis

Pseudo-aligned and pre-processed RNA-Seq data were downloaded. Samples with >70% of total genes with 0 sequence reads were deemed as very low-quality and filtered out. Normalization, batch effect correction and differential expression were performed with R package DEseq2 v1.28.1 (59). Wilcoxon rank sum test was done to determine statistical differences between categorical groups.

### Statistical analysis

Statistical analyses were carried out using R or GraphPad Prism 9 software. Shapiro-Wilk test was used to assess the normality of data distribution. Analysis of variance followed by Dunnett’s multiple comparisons test, Kruskal-Wallis test followed by post-hoc Dunn’s test or two-way ANOVA with Sidak’s multiple comparisons test were used as indicated in the figure legends. Mann-Whitney U test and Wilcoxon rank sum test were used for the analysis of unpaired and paired samples respectively. Correlations were calculated using the nonparametric Spearman correlation test. Differences between values were considered significant at p<0.05. *, p<0.05; **, p<0.01; ***, p<0.001; ****, p<0.0001. All experiments were performed at least three times in triplicate otherwise indicated.

## Author’s contributions

Conceptualization and supervision of the project: J.-P.G. and S.M. Research design: J.M.P., A.K, L.G., St.M., J.-P.G., S.M; Methodology, J.M.P., A.K, L.G., St.M., J.-P.G., S.M.; Investigation, J.M.P., A.K., V.A.G., S.H., S.M.; Data analysis, J.M.P., A.K., S.H, M.B., J.S., J.-P.G., S.M.; Reagents: St.M. ; Human samples : M.B., J.S.; Writing-original draft, J.M.P. and S.M.; Editing, J.M.P.,A.K.., St.M., J.-P.G .and S.M.; Project administration and funding, J.-P.G and S.M. All authors have read and agreed to the published version of the manuscript.

## Supporting information

Supplementary materials for Pellegrini et al 2024

## Acknowledgments

We thank Dr. P. Pierre (CIML, Marseille, France), Dr. M. Fukuda (La Jolla Cancer Research Foundation, U.S.A), Dr. M. Negroni (IBMC Strasbourg, France), Pr. S. Yamaoka (Tokyo Medical and Dental University, Japan), Pr. Ignacio Moriyón and Pr. Raquel Conde-Àlvarez for kind gifts of reagents. We acknowledge the PICSL imaging facility of the CIML (ImagImm), member of the national infrastructure France-BioImaging supported by the French National Research Agency (ANR-10-INBS-04), the cytometry facility of the CIML (S. Bigot, Dr. C. Mionnet and D. Popoff) for their kind assistance. J.M.P. is a recipient of a post-doctoral fellowship from the Fondation de Coopération Scientifique “Méditerranée Infection”.

## Funding

Work was supported by institutional funding from the Centre National de la Recherche Scientifique (CNRS) and the Institut national de la santé et de la recherche médicale (Inserm), and by the Excellence Initiative of Aix-Marseille Université (AMU) (A*MIDEX), a French “Investissements d’Avenir” programme, a French “Investissements d’Avenir” programme, by Laboratory of scientific excellence (labex) initiatives, Labex «INFORM» and «Institut Convergence CenTuri» and INSERM Transfert (CoPoC,, 2020 CITHEB), ANR-20-CE15-0016.

## Notes

### Competing Interest Statement

The authors have declared no competing interest.

