## Supplementary materials for Pellegrini et al 2024 for "SLAMF7 and SLAMF8 receptors shape human plasmacytoid dendritic cell responses to intracellular bacteria"

**This PDF file includes:**

Figures. S1 to S3

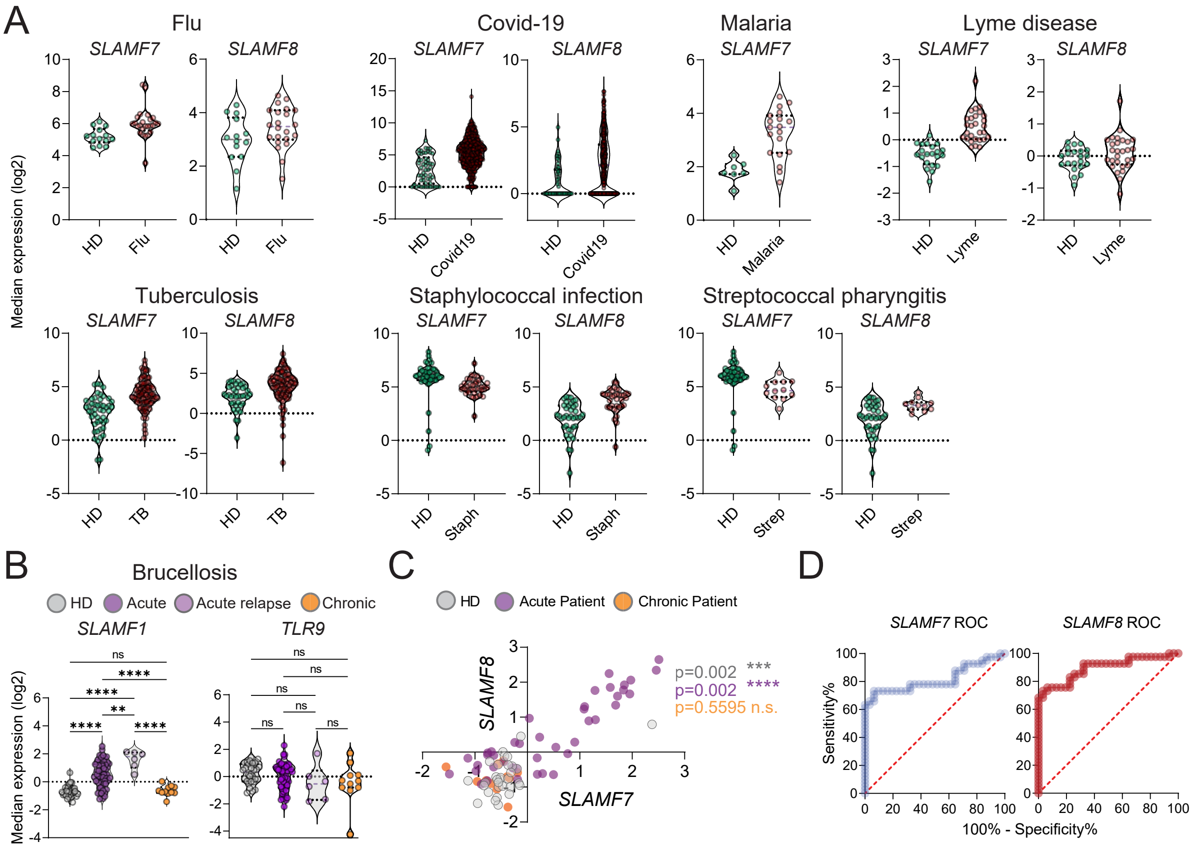


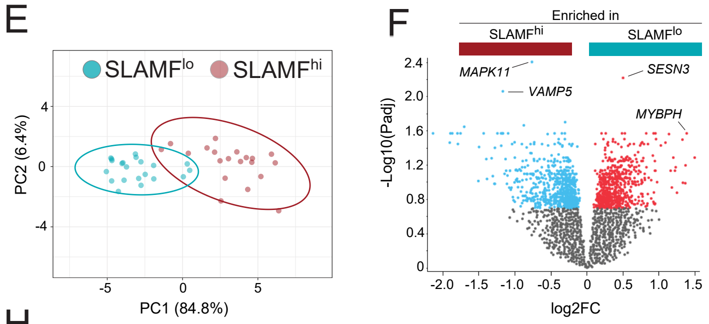


G

| Category | | SLAMF^lo | SLAMF^hi | p value |
|---|---|---|---|---|
| n | | 22 | 21 | |
| Age (years) | | 37.1 ± 4.2 | 46.0 ± 4.6 | 0.1271[a] |
| Sex | | | | |
| | Male | 19 (86.36%) | 16 (76.19%) | 0.4566[b] |
| | Female | 3 (13.64%) | 5 (23.81%) | |
| Dominant symptom | | | | |
| | General | 13 | 14 | |
| | Sacroiliitis | 0 | 1 | |
| | Epididymitis | 0 | 1 | |
| | Spondylitis | 0 | 1 | |
| | Arthritis | 6 | 0 | |
| | Arthralgia | 1 | 0 | |
| | No Data | 2 | 4 | |
| Co-morbidity | | 2 (9.09%) | 6 (28.57%) | 0.1324[b] |

H

| Category | | SLAMF^lo | SLAMF^hi | p value |
|---|---|---|---|---|
| n | | 21 | 13 | |
| Age (months) | | 18.19 ± 4.85 | 19 ± 5.3 | 0.9142[a] |
| Sex | | | | |
| | Male | 12 (57.1%) | 7 (53.8%) | >0.9999[b] |
| | Female | 9 (42.9%) | 6 (46.2%) | |
| Severity | | | | |
| | Mild | 13 (61.9%) | 9 (69.2%) | 0.7271[b] |
| | Moderate-Severe | 8 (38.1%) | 4 (30.8%) | |
| Duration symptoms (days) | | 7.6 ± 1.1 | 9.5 ± 2.0 | 0.8676[a] |
| Days past onset | | 3.1 ± 0.4 | 2.8 ± 0.5 | 0.5235[a] |
| Maximum temperature (°C) | | 37.57 ± 0.27 | 37.98 ± 0.20 | 0.2846[a] |
| Lymphocytes ($10^3$ cells/mm3) | | 5.657 ± 0.914 | 3.946 ± 0.649 | 0.1883[a] |
| Neutrophils ($10^3$ cells/mm3) | | 6.491 ± 1.112 | 7.480 ± 1.344 | 0.5794[a] |
| Monocytes ($10^3$ cells/mm3) | | 1.153 ± 0.228 | 1.048 ± 0.227 | 0.7587[a] |
| Eosinophils ($10^3$ cells/mm3) | | 0.146 ± 0.039 | 0.129 ± 0.053 | 0.7836[a] |
| Basophils ($10^3$ cells/mm3) | | 0.137 ± 0.036 | 0.073 ± 0.030 | 0.231[a] |
| Red blood cells ($10^6$ cells/mm3) | | 3.97 ± 0.15 | 3.94 ± 0.18 | 0.9005[a] |
| Platelets ($10^3$ cells/mm3) | | 308.8 ± 21.9 | 325.9 ± 289.9 | 0.6692[a] |

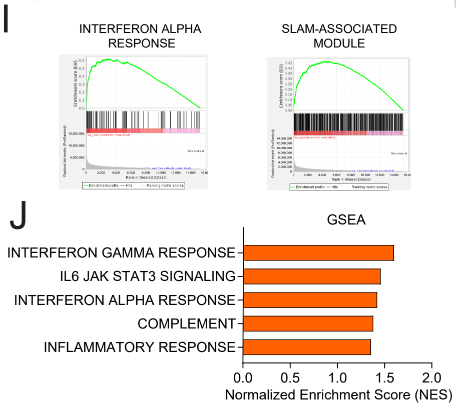


**Supplementary Figure 1. A.** Violin plots show the median expression of *SLAMF7* and *SLAMF8* normalized counts from blood transcriptomics data between healthy donors (HD, Green dots) and patients (Red dots) suffering from flu (GSE100160), Covid-19 (GSE152075), malaria (GSE116149), Lyme disease (GSE145974), tuberculosis (GSE19491), staphylococcal infection (GSE100165), and streptococcal pharyngitis (GSE158163). **B-G.** RNA seq transcriptomic profiling obtained from whole blood samples from healthy donor (HD) controls or primary brucellosis patients in acute, acute relapse or chronic phase of infection. **B.** The median expression of *SLAMF1* and *TLR9* normalized counts is shown. X-axis HD controls, n=36 (Grey); Brucellosis patients, Acute, n=54 (Purple), Acute with relapse, n=6 (Light Purple), Chronic, n=12 (Orange). Y-axis: log2 residual gene expression counts. Significant differences are shown (Multiple comparison Kruskal-Wallis test, followed by post-hoc Dunn's test). **C.** Correlation between *SLAMF7* and *SLAMF8* RNA counts across all groups of individuals including HD, acute and chronic brucellosis patients. Nonparametric Spearman correlation test. **D.** Receiver operating characteristic (ROC) curves of *SLAMF7* and *SLAMF8* gene expression level's ability to discriminate acute from chronic brucellosis. **E.** PCA analysis of blood signature in $SLAMF^{hi}$ and $SLAMF^{lo}$ categorized brucellosis patients. **F.** Volcano plot showing differentially expressed genes between $SLAMF^{hi}$ and $SLAMF^{lo}$ brucellosis patients. (Red: enriched in $SLAMF^{lo}$ patients, Light Blue: enriched in $SLAMF^{hi}$ patients). **G.** Demographic and clinical data of $SLAMF^{hi}$ and $SLAMF^{lo}$ categorized brucellosis patients**.** Continuous data are expressed as mean ± SD, and categorical data are expressed as number (percentages). p values were calculated by (a) Mann-Whitney U test for unpaired and non-parametric samples and by (b) Fisher exact test for categorical variables. **H.** Demographic and clinical data of $SLAMF^{hi}$ and $SLAMF^{lo}$ categorized salmonellosis patients from the second cohort (GSE69529). Continuous data are expressed as mean ± SEM, and categorical data are expressed as number (percentages). p values were calculated by (a) Mann-Whitney U test for unpaired and non-parametric samples and by (b) Fisher exact test for categorical variables. **I, J.** Gene set enrichment analysis (GSEA) was performed on RNA-Seq data from $SLAMF^{hi}$ and $SLAMF^{lo}$ categorized salmonellosis patients (GSE69529). **I.** Individual gene set enrichment plots for Interferon alpha response and SLAM-associated module are shown. J. Top five gene sets according to the normalized enrichment score (NES). FDR q-val<0.05.

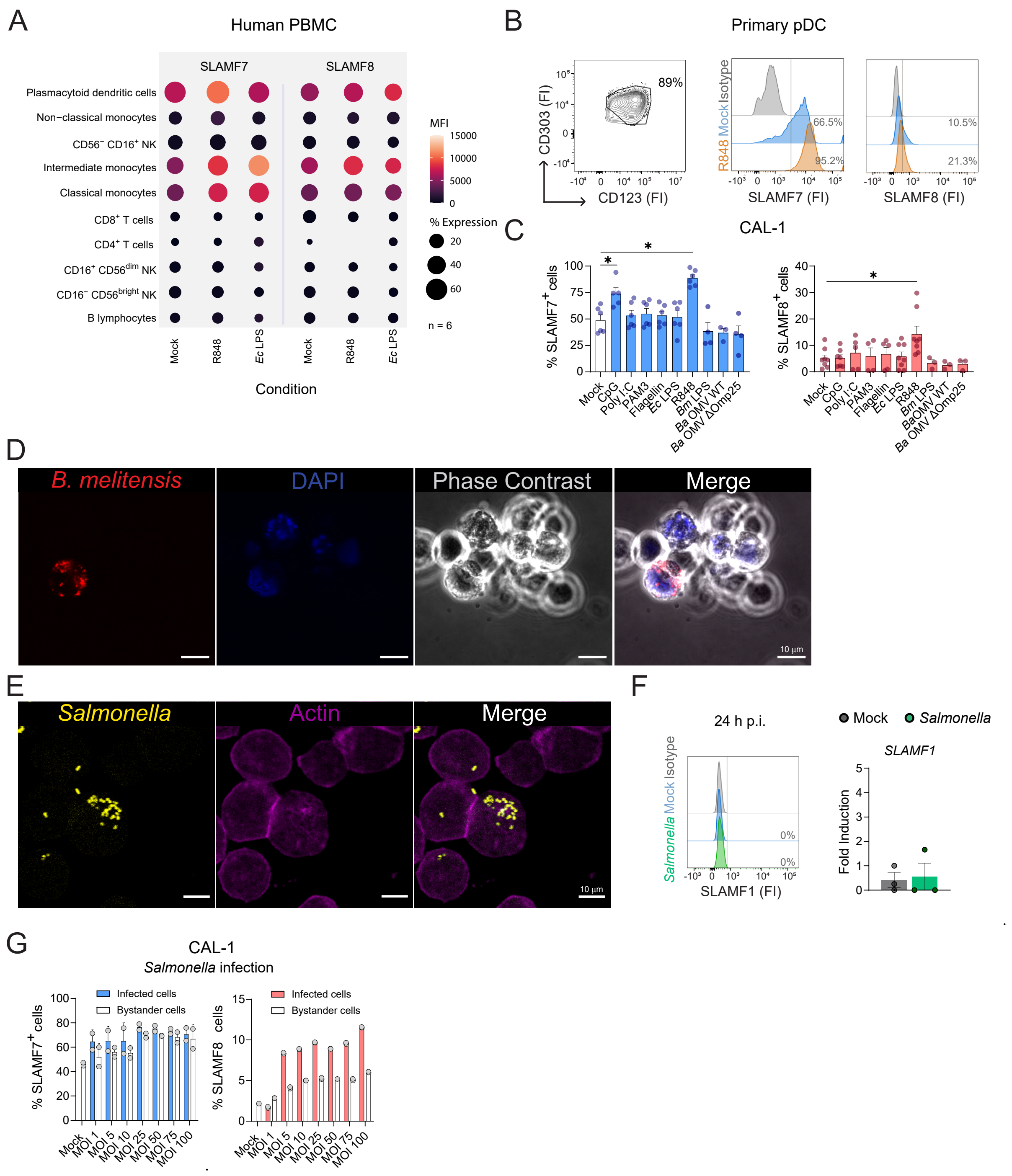


**Supplementary Figure 2**. **A.** SLAMF7 and SLAMF8 surface expression in different cell types from peripheral blood mononuclear cells (n=6) from Healthy donors (HD), treated or not with R848 or *E. coli* LPS (100 ng/mL) for 24 h, and analyzed by spectral flow cytometry. Dot plots represent the median fluorescence intensity (MFI, color) and the percentage of expression in a determined cell type (size). Color and size codes are indicated on the right. **B.** Human plasmacytoid dendritic cells (pDC) were purified from healthy individuals' blood. pDC purity was analyzed by flow cytometry based on the expression of CD303 and CD123 (left panel). Then, cells were stimulated (Orange) or not (Blue) with R848 (100 ng/mL) for 24 h, and SLAMF7 and SLAMF8 expression was evaluated by flow cytometry. Representative histograms of flow cytometry experiments showing SLAMF7 (middle) and SLAMF8 expression (right) are shown. **C.** CAL-1 cells were stimulated with CpG (TLR9 ligand, 100 ng/mL), PolyI:C (TLR3 ligand, 100 ng/mL), PAM3CSK4 (TLR1/2 ligand, 100 ng/mL), flagellin (TLR5 ligand, 100 µg/mL), *E. coli* LPS (TLR4 ligand, 100 ng/mL), R848 (TLR7/8 ligand, 100 ng/mL), *B. melitensis* LPS (10 ng/mL), WT *B. abortus* outer membrane vesicles (10 µg/mL), or ΔOmp25 *B. abortus* outer membrane vesicles (10 µg/mL) for 24 h. Then, SLAMF7 and SLAMF8 expression was evaluated by flow cytometry. Column graphs showing the percentage of SLAMF7$^+$ (left, Blue) and SLAM/8$^+$ (right, Pink) cells. Mean ± SD. n=8. Significant differences are indicated. Statistical differences were all calculated using One-way ANOVA followed by Dunnett's multiple comparisons test. *, $p < 0.05$. **D.** Representative confocal images showing CAL-1 cells infected with *Brucella melitensis* (M.O.I. of 5000) at 48 h post infection (p.i.). Scale bars: 10 µm. n=3. **E.** Representative confocal images of CAL-1 infected with DsRed WT *Salmonella* Typhimurium at 24 h p.i.. *Salmonella* (Yellow) and Actin (Purple) are shown. Scale bars: 10 µm. n=3. **F.** CAL-1 cells were infected with WT *S.* Typhimurium (M.O.I. of 25, Green) or not (Grey) for 24 h. Then, SLAMF1 expression were evaluated by flow cytometry and RT-qPCR. n=3. Left: representative histograms of flow cytometry experiments are shown. Right: Column graphs showing *SLAMF1* gene expression relative to housekeeping gene, *ACTB*. Mean ± SEM. **G.** CAL-1 cells were infected with DsRed WT *S.* Typhimurium at the indicated M.O.I. for 24 h. Then, SLAMF7 and SLAMF8 expression was evaluated by flow cytometry on infected (filled colored bars; SLAMF7$^+$, Blue; SLAMF8$^+$, Pink) and bystander cells (empty bars). Column graphs showing the percentage of SLAMF7$^+$ and SLAMF8$^+$ cells. Mean ± SD. n=3. Statistical differences were all calculated using One-way ANOVA followed by Dunnett's multiple comparisons test. no p value, non-significant.

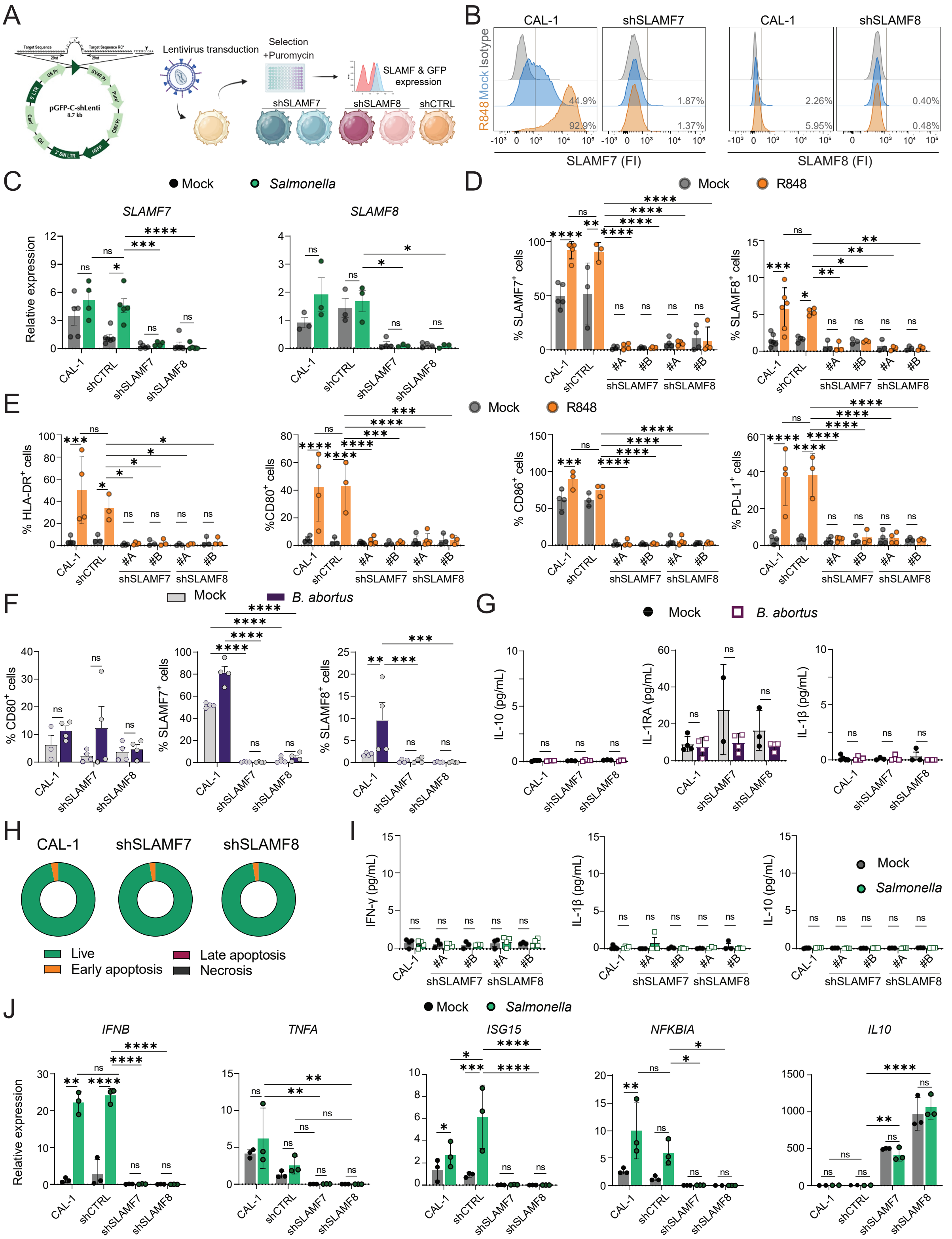


**Supplementary Figure 3**. **A.** Strategy for generating and selecting stable specific SLAMF7 and SLAMF8-knockdown (KD) CAL-1 cells using lentivirus transduction. **B.** WT CAL-1 cells, non-targeting shRNA-transduced control cells (shCTRL) and SLAMF-silenced cells (shSLAMF7 and shSLAMF8) were stimulated with the TLR7 ligand R848 (100 ng/mL) for 24 h. Representative histograms showing SLAMF7 and SLAMF8 surface expression demonstrate efficient SLAMF7- or SLAMF8- silencing in CAL-1 cells at resting state (Mock, Blue) and upon stimulation (R848, Orange). **C.** CAL-1 and SLAMF-silenced cells were infected with DsRed WT *Salmonella* Typhimurium (M.O.I. of 25, Green) for 3 h. SLAMF7 (left) and SLAMF8 (right) gene expression was evaluated by RT-qPCR. Column graphs showing relative gene expression to housekeeping gene, *ACTB*. Mean ± SD. n=3-5. **D, E.** WT CAL-1 cells, non-targeting shRNA-transduced control cells (shCTRL) and SLAMF-silenced cells (shSLAMF7 and shSLAMF8, 2 independent clones, A and B, analyzed per type of SLAM-KD cells) were stimulated or not with the TLR7 ligand R848 (100 ng/mL, orange) for 24 h. Expression of SLAMF7 (D, left) and SLAMF8 (D, right) and HLA-DR, CD80, CD86 and PD-L1 surface markers (E, from left to right) were determined by flow cytometry. Column graphs showing the percentage of positive cells. Mean ± SD. n=4-5. **F, G.** WT CAL-1 and SLAMF-KD cells were infected with mCherry WT *Brucella abortus* or mCherry WT *B. melitensis* (M.O.I. of 5000, Purple) for 48 h. **F.** Surface expression of CD80, SLAMF7 and SLAMF8 was determined by flow cytometry. Column graphs showing the percentage of positive cells. Mean ± SD. n=4. **G.** Cytokine secretion (pg/ml) was determined in culture supernatants (from **F**) by multiplex assay and shown by column graphs. Mean ± SD. n=3-4. **H-J.** WT CAL-1 and SLAMF-silenced cells were infected with DsRed WT *Salmonella* Typhimurium (M.O.I. of 25) for 24 h (H, I) or 3 h (J). **H.** No change in viability was observed in WT or silenced CAL-1 cells. Cells were subjected to dual staining with annexin V (Ann V) and propidium iodide (PI), and cell death was assessed by flow cytometry. Parts of the whole represent the percentage of live cells (Green) or cells in early apoptosis (Ann $V^+$ $PI^-$, Orange), late apoptosis (Ann $V^+$ $PI^+$, Pink) or necrosis (Ann $V^-$ $PI^+$, Purple). **I.** No IFN-$\gamma$, IL-1$\beta$ or IL-10 secretion was triggered in human pDC by *S*. Thyphimurium infection. Cytokine secretion was determined in culture supernatants using multiplex assay. Column graphs showing cytokine concentration (pg/ml). Mean ± SD. n=4. **J.** *IFNB*, *TNFA*, *ISG15*, *NFKBIA*, and *IL10* gene expression was evaluated by RT-qPCR. Column graphs showing relative gene expression to housekeeping gene, *ACTB*. Mean ± SD. n=3. Statistical differences were all calculated using Two-way ANOVA followed by Sidak's multiple comparisons test. *, p < 0.05; **, p < 0.01; ***, p < 0.001, ****; p < 0.0001. ns., non-significant.
